# A genotype independent *DMP*-HI system in dicot crops

**DOI:** 10.1101/2021.06.21.449224

**Authors:** Yu Zhong, Baojian Chen, Dong Wang, Xijian Zhu, Yuwen Wang, Mengran Li, Yifan Li, Jinchu Liu, Jinzhe Zhang, Ming Chen, Min Wang, Tjitske Riksen, Xiaolong Qi, Dehe Cheng, Zongkai Liu, Jinlong Li, Chen Chen, Yanyan Jiao, Wenxin Liu, Bin Yi, Sanwen Huang, Chenxu Liu, Kim Boutilier, Shaojiang Chen

## Abstract

Doubled haploid (DH) technology is used to obtain homozygous lines in a single generation, which significantly accelerates the crop breeding trajectory. Traditionally, *in vitro* culture is used to generate DHs, but is limited by species and genotype recalcitrance. *In vivo* haploid induction (HI) through seed is been widely and efficiently used in maize and was recently extended to several monocot crops. However, a similar generic and efficient HI system is still lacking in dicot crops. Here we show that genotype-independent *in vivo* HI can be triggered by mutation of *DMP* genes in tomato, rapeseed and tobacco with HI rates of ~1.9%, 2.4% and 1.2%, respectively. The *DMP*-HI system offers a robust DH technology to facilitate variety improvement in these crops. The success of this approach and the conservation of *DMP* genes paves the way for a generic and efficient genotype-independent HI system in other dicot crops.

## INTRODUCTION

The rapid development of high yield and quality crop varieties is essential to ensure world-wide food and commodity security. One of the most important aspects of crop breeding is the development of homozygous lines. Homozygous lines can be developed by repeated rounds of selfing or backcrossing, usually four to six generations depending on the desired level of homozygosity, which is a costly and time-consuming process (Heusden and Lindhout, 2018). Alternatively, homozygous lines can also be obtained in a single generation by using DH technology (Jacquier et al., 2020). Haploid embryos can be induced *in vitro* from cells of the male or female gametophyte or *in vivo* in seeds by interspecific crosses or by intraspecific crosses with haploid inducer lines (Kalinowska et al., 2019; Lv et al., 2020; Wang et al., 2021). Of these methods, *in vivo* HI triggered by inducer lines is the most efficient approach, but it is currently limited to few monocot crops. Although HI systems have been reported for some dicot crops, unlike monocot crops, these systems are either inefficient or have not been extended to other dicot crops (Hougas and Peloquin, 1957; Hougas et al., 1958; Fu et al., 2018; Jacquier et al., 2020; Hooghvorst and Nogués, 2020a). Dicots account for the majority of the angiosperm species, including many economically important vegetable, fruit, seed and industrial crops like tomato (*Solanum lycopersicum*), chili pepper (*Capsicum annuum*), cucumber (*Cucumis sativus*), rapeseed (*Brassica napus*), soybean (*Glycine max*), cotton (*Gossypium hirsutum*) and tobacco (*Nicotiana tabacum*). However, many important dicot crops are completely recalcitrant for haploid induction, e.g., tomato and cotton, while almost all crops have recalcitrant genotypes (Jacquier et al., 2020; Hooghvorst and Nogués, 2020a; Hooghvorst and Nogués, 2020b). The lack of suitable HI systems for many dicot crops means that the slower and more costly classical breeding approach is still required in the development of homozygous lines.

Significant breakthroughs in DH production have been made in the last few years through the identification of genes that induce maternal haploid embryos *in vivo* in maize. Maternal HI systems rely on pollination by specific male inducer lines that stimulate the haploid egg cell to develop into an embryo that lacks the parental chromosome component (Hougas and Peloquin, 1957; Hougas et al., 1958; Hussain and Franks, 2019; Jacquier et al., 2020). Naturally occurring HI lines have been used extensively in maize breeding programs since the 1950s (Coe, 1959), and this trait was rapidly engineered in a number of monocot crops after identification of the *ZmPHOSPHOLIPASE-A1 /MATRILINEAL/NOT LIKE DAD (ZmPLA1 /MTL/NLD*) HI gene (Kelliher et al., 2017; Liu et al., 2017; Gilles et al., 2017; Yao et al., 2018; Zhong et al., 2019; Liu et al., 2019; Liu et al., 2020).

*ZmPLA1/MTL/NLD* genes have only been identified in monocots (Kelliher et al., 2017; Yao et al., 2018; Liu et al., 2019), while genes related to a second maize HI gene, *Zea may DOMAIN OF UNKNOWN FUNCTION 679 membrane protein (ZmDMP*), have been identified in both monocots and dicots (Zhong et al., 2020). The utility of *dmp* mutants for HI in dicots was recently demonstrated in the model plant arabidopsis (*Arabidopsis thaliana*) (Zhong et al., 2020), but it is not known whether this approach can be translated to dicot crops. Here, we developed a method to identify candidate *DMP* genes for HI in crops and demonstrate *DMP*-mediated maternal HI in three major dicot crops with different ploidy levels and from two different plant families, tomato, rapeseed and tobacco. This breakthrough, together with genotype independent HI in tomato, provides proof-of-concept for the development of a universal *DMP*-HI system in dicot crops.

## RESULTS

### *DMP* genes in dicot crops

Our previous results identified *ZmDMP* orthologues in multiple species, but with an average amino acid sequence identity of 66%. Moreover 42% of dicots contain multiple *DMP* gene copies as a result of genome duplication and/or interspecific hybridization (Zhong et al., 2020). The relatively low sequence identity and the presence of multiple gene copies makes it difficult to accurately identify *ZmDMP* orthologues for the development of a *DMP*-HI system in dicot crops. To this end, DMP proteins from seven dicot crops with the highest amino acid sequence identity with ZmDMP were each used as a query to search the corresponding genome database of each crop. *DMP* genes with >50% amino acid identity were selected for expression analysis using public transcriptome databases. We identified: up to four *DMP* genes in rapeseed (*B. napus) (BnDMP1A/BnaA03g55920D; BnDMP1C/BnaC03g03890D*; *BnDMP2A/BnaA04g09480D; BnDMP2C/BnaC04g31700D*) that are all highly expressed in anthers and flower buds; a single *DMP* gene in tomato (*Solanum lycopersicon*) (*SlDMP*/Solyc05g007920) that is highly expressed in pollen and flower buds (Zhong et al., 2020); a single *DMP* gene in chili pepper (*Capsicum annuum*) (*CaDMP*/*Capana04g002148*) that is highly expressed in closed flower bud and open flower; two cotton (*Gossypium hirsutum*) *DMP* genes (*GhDMP1*/LOC107911807; *GhDMP2*/LOC107924398) that are expressed in the stamen; two soybean (*Glycine max*) *DMP* genes (*GmDMP1/GLYMA_18G097400; GmDMP2/GLYMA_18G098300*) expressed in flower bud; and a single cucumber (*Cucumis sativus*) *DMP* gene (*CsDMP/Csa_1G267250*) that is expressed in male flower bud (Supplemental Table 1). Expression data was not available for tobacco (*Nicotiana tabacum*), which contained three *DMP* genes (*NtDMP1* /LOC107762412; *NtDMP2.*/LOC107783066; *NtDMP3*/LOC107807404). Like DMP proteins in maize and arabidopsis (Takahashi et al., 2018; Cyprys et al., 2019; Zhong et al., 2019; Zhong et al., 2020), all of the above DMP proteins have a DUF679 domain and multiple transmembrane (TM) helices (Supplemental Table 1). Alignment of these DMP proteins showed that the entire DUF679 domain (>56% identity) and especially the first predicted transmembrane helices (>80% identity) are conserved among these species (Supplemental Figure 1 and Supplemental Table 2).

Next, a complementation strategy was used to determine whether these candidate *DMP* genes have potential HI functions, as measured by their ability to rescue the reduced seed set phenotype of the arabidopsis *dmp8dmp9* HI mutant. To this end, eight of the above *DMP* genes were expressed with the *AtDMP9* promoter in the *dmp8dmp9* background, and seed setting evaluated in T1 plants. All of the eight genes significantly increased the seed set of the arabidopsis *dmp8dmp9* mutant (Figure 1), indicating that these *DMP* genes can complement *AtDMP8* and *AtDMP9* functions during double fertilization (Takahashi et al., 2018; Cyprys et al., 2019; Zhong et al., 2020), and could be used to develop a HI system in these dicots. Tomato, rapeseed and tobacco have well-established genetic transformation systems and are relatively easy to hybridize, which led us to explore the possibility of developing *DMP*-HI systems in these three crops.

**Figure 1.**
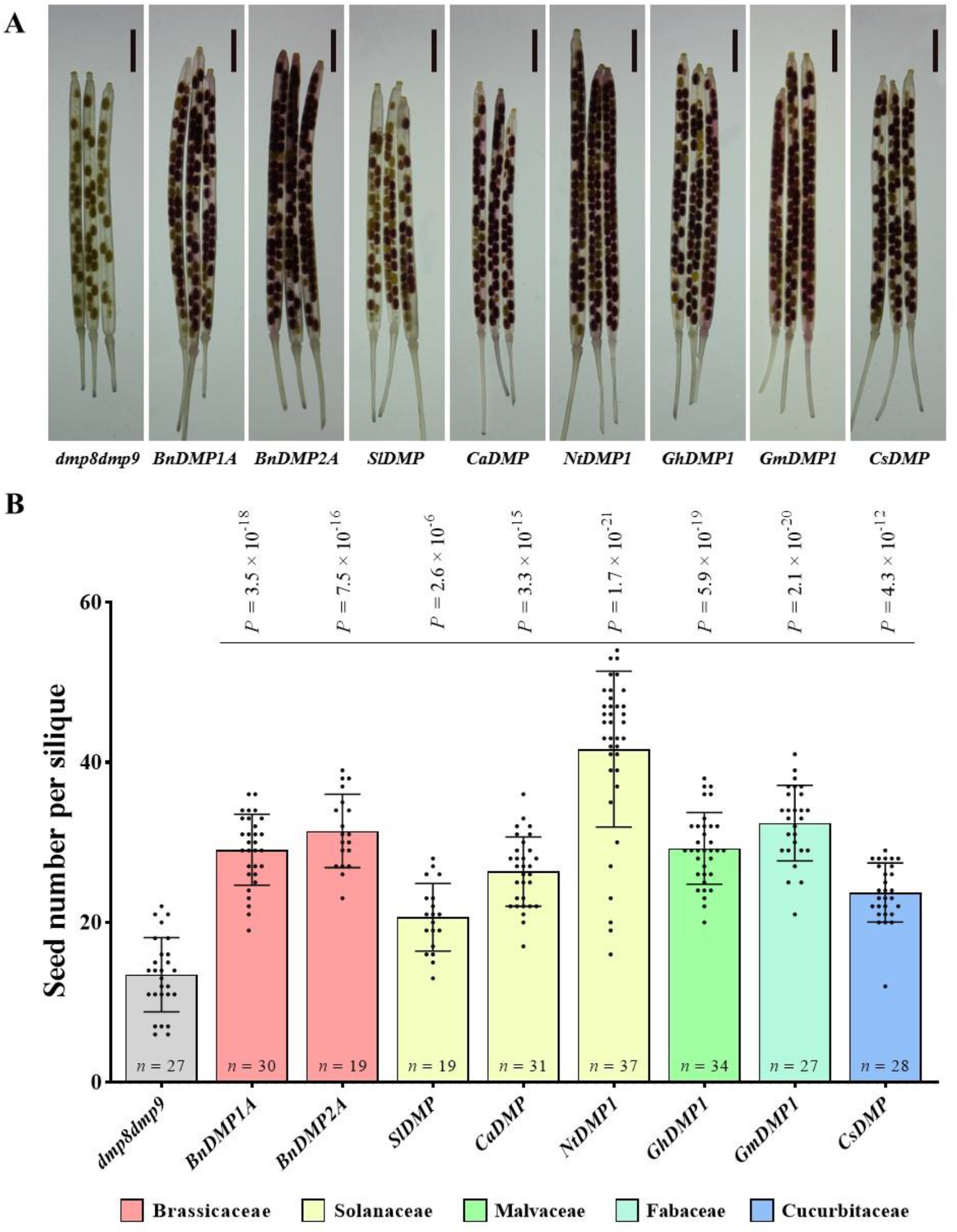
*DMP* genes from multiple dicot crops complement *dmp8dmp9* phenotypes in arabidopsis. (**A**) Representative images of siliques from the *dmp8dmp9* mutant and different complementation lines in the *dmp8dmp9* background. Scale bar, 2 mm. (**B**) Quantification of seed number per silique in the *dmp8dmp9* mutant and the corresponding complementation lines. The bars with different colors indicate different plant families. Data represent the mean ± s.d.; \*\*\**p* < 0.001 (two-tailed Student’s *t*-test); *n*, number of siliques. The detailed information of each gene are described in Supplemental Table 1.

### Haploid seed induction, identification, and DH production in tomato

A CRISPR-Cas9 mutagenesis construct was designed to generate *dmp* loss-of-function mutants in the three tomato cultivars (Figure 2A). The construct also includes the FAST-Red marker for haploid identification (Zhong et al., 2020). Mutants with insertions and/or deletions that resulted in translational frame shifts and premature stop codons were found in the T0 generation of the Ailsa Craig, Micro-Tom and Moneyberg genotypes (Supplemental Table 3). Homozygous or biallelic *sldmp* mutants were chosen for subsequent experiments (Figure 2B). Pleiotropic seed phenotypes were observed in self-pollinated fruits from these *sldmp* mutants (Figure 2C-2E and Supplemental Figure 2 and Supplemental Figure 3A-3C and Supplemental Figure 4A-4C). Compared to wild type plants, the number of filled seeds was significantly reduced (Figure 2D and Supplemental Figure 3B and Supplemental Figure 4B). and the percentage of both aborted seeds and undeveloped ovules significantly increased (Figure 2E and Supplemental Figure 3C and Supplemental Figure 4C) in *sldmp* mutants. Reciprocal crosses between wild type and the Ailsa Craig *dmp* mutant and analysis of pollen germination showed that the abnormal seed phenotypes are due to a paternal (pollen) fertilization defect (Supplemental Figure 5), as previously shown for arabidopsis (Takahashi et al., 2018; Cyprys et al., 2019).

**Figure 2.**
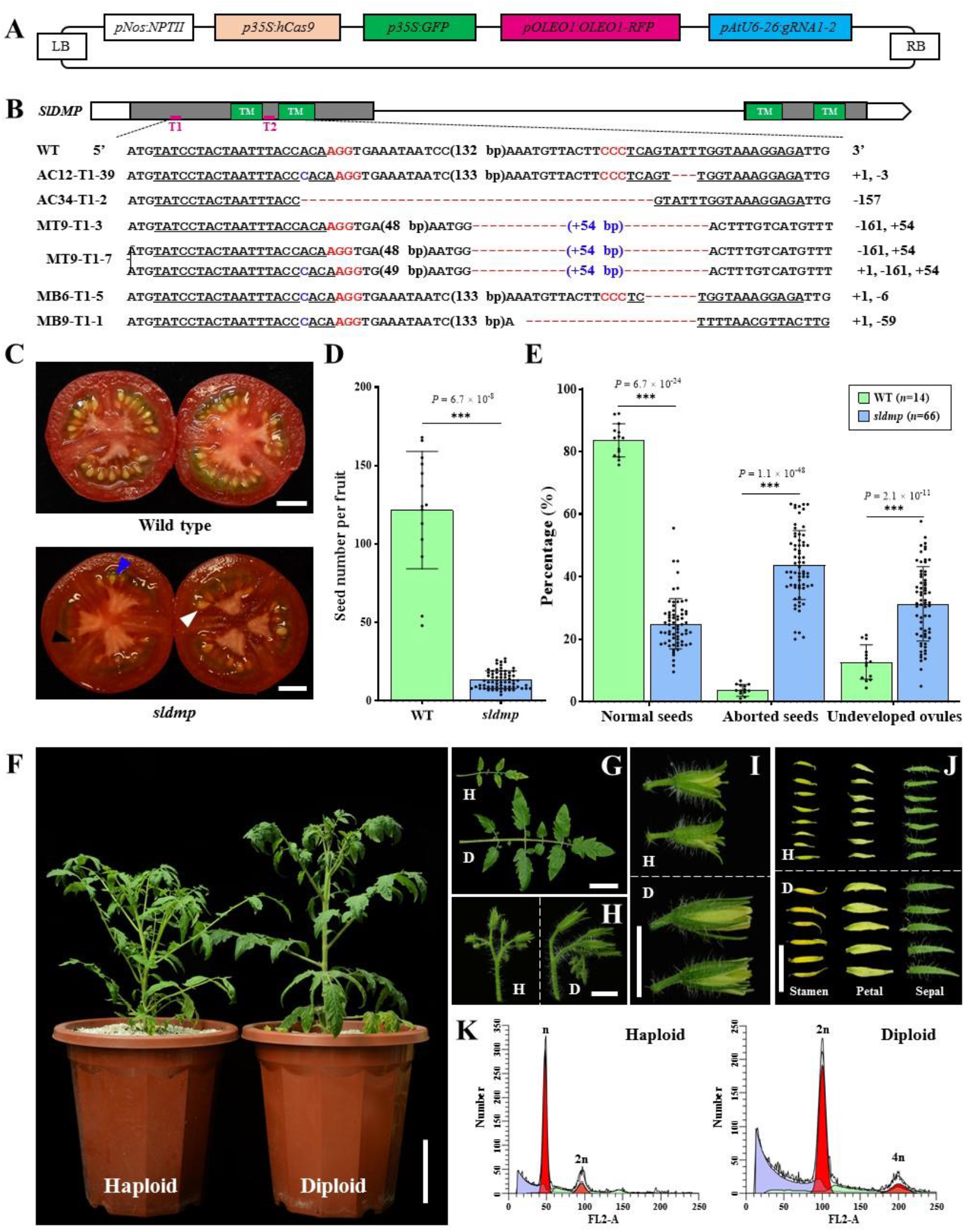
Mutation of tomato *SlDMP* induces haploids. (**A**) The CRISPR/Cas9 mutagenesis vector comprising two sgRNAs (*gRNA1-2*) targeting *SlDMP,* and the *pNos:NPTII* and *p35S:GFP* and FAST-Red selection cassettes. (**B**) Schematic representation of the wild-type (WT) *SlDMP* gene. Filled blocks, clear blocks and the gray line indicate the coding region, the untranslated regions, and the intron, respectively. Green blocks correspond to the four predicted transmembrane domains (TM). Pink lines indicate the two regions (T1, T2) targeted by the sgRNAs. The sequences from wild type (WT) and mutant alleles from three backgrounds (AC, Ailsa Craig; MT, Micro-Tom; MB, Moneyberg) are shown below the overview. The sgRNA target sequences are underlined, and the protospacer-adjacent motif (PAM) is shown in red. Nucleotide insertions are shown in blue and deletions by red dashes. (**C**) Representative images of ripe fruit from selfed WT and CRISPR-Cas9 *sldmp* mutants in the Ailsa Craig background. White, black, and blue arrowheads indicate normal seeds, aborted seeds and undeveloped ovules, respectively. (**D** and **E**) Quantification of seed number (D) and seed phenotypes (E) in fruits shown in (C). Data represent the mean ± s.d.; \*\*\**p* < 0.001 (two-tailed Student’s *t*-test); *n*, number of fruits. (**F** to **J**) Phenotypes of plants (F), leaves (G), inflorescences (H), flower buds (I) and dissected flower parts (J) of haploid (H) and diploid (D) plants. (**K**) Flow cytometry verification of the ploidy of a putative haploid and a diploid control. The *x* axis represents the signal peak for the nucleus and the *y* axis represents the number of nuclei. Scale bars: 1 cm (C, H, I and J), 10cm (F) and 5 cm (G). In (C) and (K), experiments were repeated at least three times and similar results were obtained.

To determine whether *sldmp* mutants can induce haploids upon selfing, we sowed selfed seeds from T1 progenies of *sldmp* mutants in the Ailsa Craig background. In the absence of segregating molecular markers, we first identified putative haploid plants based on their phenotype, i.e. smaller organs and sterility (Ravi and Chan, 2010; Zhong et al., 2019; Zhong et al., 2020). Among 55 T1 plants, one plant (1.8%), which was relatively shorter and bushier than the wild-type control (Figure 2F), showed the typical haploid phenotype (Figure 2G-2J). This plant was subsequently confirmed by ploidy analysis to be a true haploid (Figure 2K). Given the low frequency of spontaneous haploid seedling production in tomato (from 9 × 10^-5^ to 4 × 10^-4^) (Cook, 1936; Koornneef et al., 1989; Hamza et al., 1993), our data suggests that mutation of the tomato pollen-expressed *DMP* gene facilitates *in vivo* haploid embryo development.

Next, we crossed a range of wild-type female plants (Supplemental Table 4) from different genetic backgrounds with *sldmp* mutants in the Ailsa Craig, Micro-Tom and Moneyberg backgrounds to determine whether *sldmp* pollen can also induce maternal haploids upon outcrossing. All crosses showed the reduced seed set and abnormal ovule/seed phenotypes observed in the wild-type × *sldmp* crosses (Figure 3A-3C and Supplemental Figure 2 and Supplemental Figure 3D-3F and Supplemental Figure 4D-4F). Nineteen haploids derived from eight different backgrounds were first screened by molecular markers and then confirmed by flow cytometry and plant phenotype (Supplemental Figure 3G-3I and Supplemental Figure 4G-4I and Supplemental Table 5). To further confirm the maternal origin of these haploids, three of these haploid seedlings were used for whole-genome resequencing. Chromosome dosage and single nucleotide polymorphism (SNP) analysis showed that none of the seedlings was aneuploid or carried paternally-derived SNPs, suggesting that *sldmp* induces ‘clean’ maternal haploids i.e. haploids lacking any paternal genome fragments (Supplemental Figure 6 and Supplemental Table 6). These results confirmed that *sldmp* mutants in one genotype can be used for clean maternal haploid induction in the same or a different genotype.

**Figure 3.**
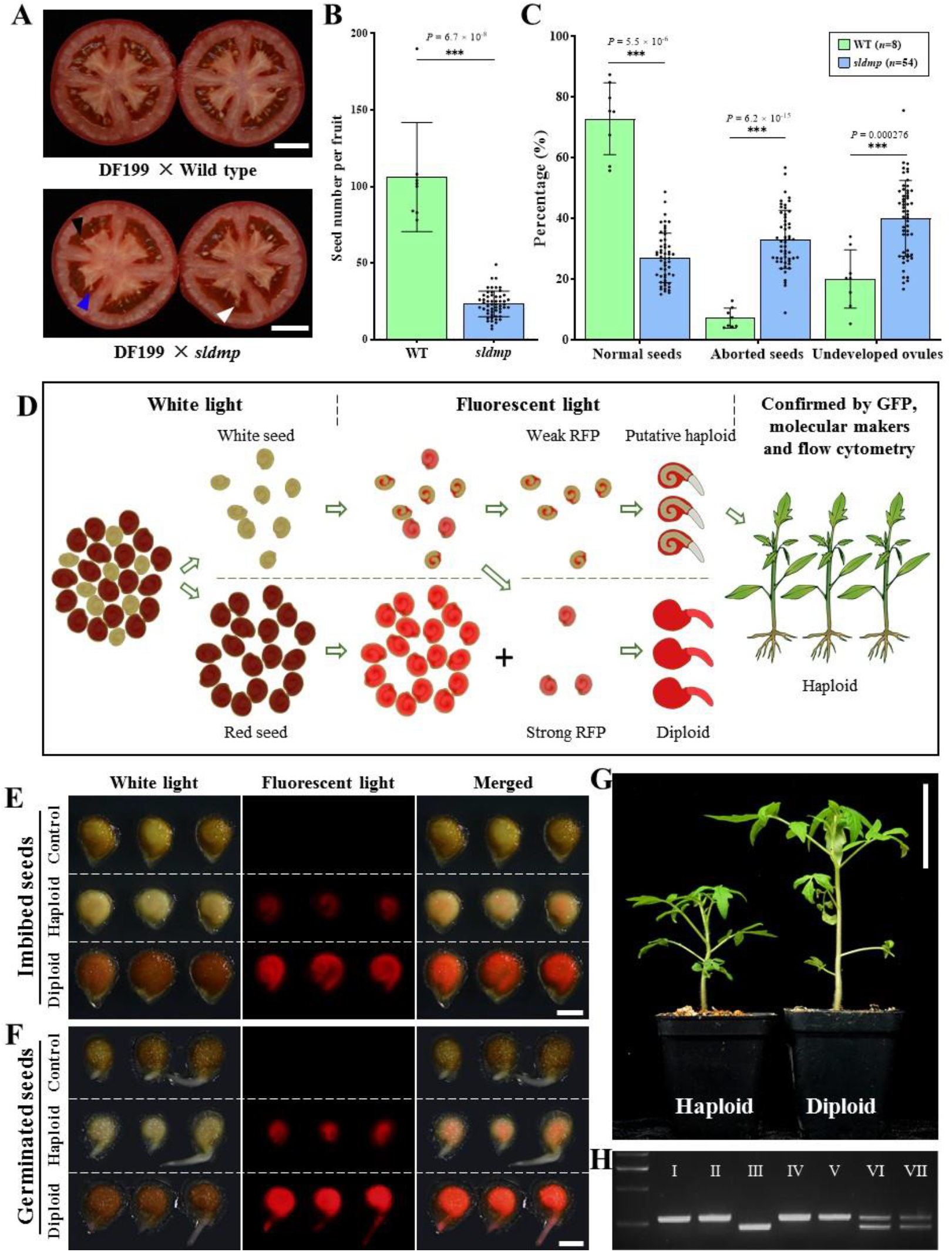
Haploid production through *sldmp* outcrossing and FAST-Red-based haploid seed identification. (**A**) Representative images of DF199 ripened fruit pollinated by wild type and an *sldmp* mutant in the Ailsa Craig background. White, black, and blue arrowhead indicate normal seeds, aborted seeds, and undeveloped ovules, respectively. (**B** and **C**) Quantification of seed number (B) and seed phenotypes (C) in fruits derived after DF199 pollination by wild type and *sldmp* mutant pollen. Data represent the mean ± s.d.; \*\*\**p* < 0.001 (two-tailed Student’s *t*-test); *n*, number of fruits. WT, wild type. (**D**) Schematic overview of haploid identification using the FAST-Red marker in the *sldmp* inducer line. After NaOCl treatment, white and red seeds can be easily distinguished under white light. Under fluorescent light, a few white seeds with strong RFP expression were regrouped. During germination, both groups were further checked for the absence/presence of RFP in the root tip. Seeds with weak RFP expression do not show RFP signal in embryo root tip and are putative haploids. These putative haploids can be further confirmed by molecular markers and ploidy analysis. (**E** and **F**) FAST-Red-based haploid seed identification. White light (left panel), fluorescent light (middle panel) and merged (right panel) micrographs of control, haploid and diploid seeds in the imbibed (E) and germinated (F) state. Control seeds were derived from a DF199 × Ailsa Craig cross. Haploid and diploid seeds were derived from DF199 × *sldmp* (Ailsa Craig) cross. (**G**) Representative images of tomato haploid and diploid seedlings. (**H**) Seedlings from putative haploids were genotyped with polymorphic markers between the inducer line and testers. The left lane shows the DNA size marker, and the Roman numbers I to VII represent the PCR products in DF199 (I), AF01 (II), *sldmp* mutant in Ailsa Craig background (III), haploid from DF199 × *sldmp* (IV), haploid from AF01 × *sldmp* (V), diploid from DF199 × *sldmp* (VI) and diploid from AF01 × *sldmp* (VII). Scale bars: 1 cm (A), 2 mm (E and F) and 5 cm (G). In A, E, F, G and H, experiments were repeated at least three times and similar results were obtained.

The arabidopsis Fast-Red marker has been used as a simple and efficient method to facilitate high throughput identification of haploids from *dmp* crosses (Zhong et al., 2020). To this end, Fast-Red marker expression was evaluated in wild-type seeds and seeds from two selfed *sldmp* mutant lines carrying the homozygous Fast-Red marker. Red/RFP-positive seeds were observed among the imbibed *sldmp* seeds under white light/fluorescent light, but only white/RFP-negative seeds were observed among the imbibed wild-type seeds (Supplemental Figure 7A). Next we separated the *sldmp* and wild-type seeds into the embryo, endosperm and seed coat components. *sldmp* embryos and endosperm were red/RFP-positive under white/fluorescent light, with the endosperm showing a weaker red color/RFP expression than the embryo, while none of the wild-type seed components were red/showed RFP expression (Supplemental Figure 7B). In line with our observations in imbibed seeds, red color/RFP were also observed under white light/fluorescent light in root tips of *sldmp* embryos during germination, but not in the root tips of the germinating wild-type embryos (Supplemental Figure 7C). These data show that the FAST-Red reporter is reliably expressed in the embryo and endosperm of tomato seeds.

To determine whether the Fast-Red marker can be used to identify tomato haploids in *dmp* crosses, we analyzed Fast-Red expression in seeds from a cross between a female wild-type DF199 line and a male *dmp* Ailsa Craig line. Imbibed seeds were first classified into red and white seed groups based on their color under white light (Figure 3, D and E). Under fluorescent light, the red seeds showed weak RFP expression in the endosperm and strong RFP expression in the embryo, while the majority of the white seeds also showed weak RFP expression in the endosperm but lacked RFP expression in the embryo (Figure 3D and 3E and Supplemental Figure 8). Some of the seeds that were initially scored as white under white light showed RFP expression in the embryo and endosperm under florescent light and were recategorized as red/RFP-expressing seeds. These two groups were further confirmed by checking root tip RFP expression during germination (Figure 3D and 3F). The red seeds with RFP expression in the embryo and endosperm were considered to carry diploid embryos, while the white seeds with weak RFP in the endosperm and no RFP expression in the embryo were considered to have maternal haploid embryos that developed spontaneously in the absence of fertilization or without the paternal chromosome component. To test this hypothesis, we sowed 218 putative haploid seeds that only showed weak RFP expression in the endosperm and 2303 putative diploid seeds that showed strong RFP expression in both the embryo and endosperm and confirmed their ploidy by molecular marker and ploidy analysis at the seedling stage (Figure 3G and 3H). We showed that all of the putative haploids were true haploids, and that all of the putative diploids were true diploids i.e., that there were no false positives or false negatives in the two seed groups (Supplemental Table 7). The FAST-Red selection procedure outlined above can therefore be used with 100% accuracy for identification of maternal haploids in tomato.

Next, we used the Fast-Red marker for haploid seed selection in crosses between diverse female tomato genotypes (Supplemental Table 4) and male *sldmp* FAST-Red lines. FAST-Red expression was stable among multiple female backgrounds after outcrossing with *sldmp* FAST-Red lines in the Ailsa Craig background (Supplemental Figure 9). The haploid induction rate (HIR) after crossing 36 different female genotypes with the *sldmp* inducer lines ranged from 0.5% to 3.7%, with an average HIR of 1.9% (Table 1). These data together with haploid induction in an additional six crosses (Supplemental Table 5) indicate that *sldmp* mutants in a given genotype can be used for efficient maternal haploid induction in the same or different genotype.

**Table 1.**
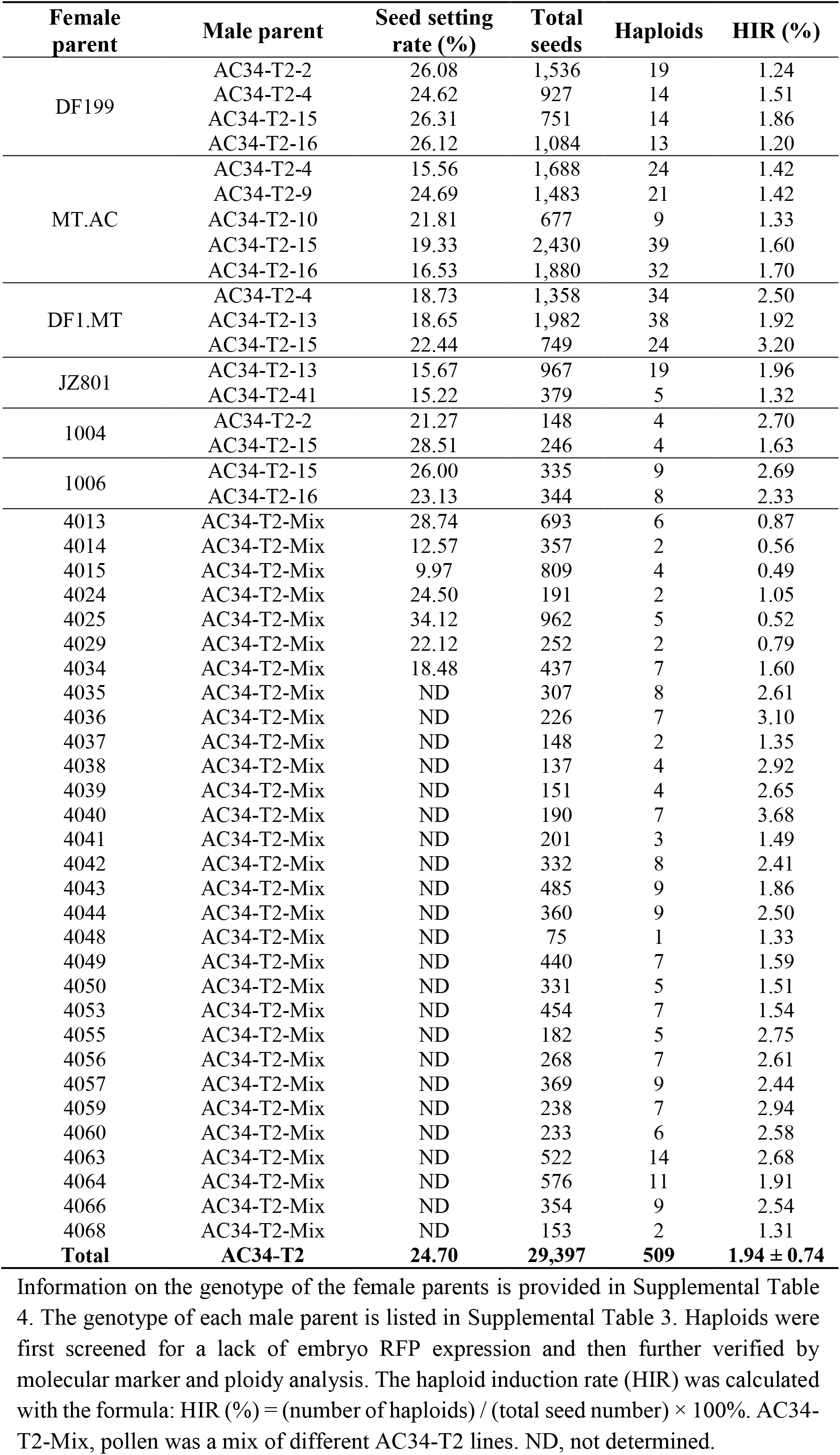
Haploid induction rate in *sldmp* mutant crosses.

| **Female parent** | **Male parent** | **Seed setting rate (%)** | **Total seeds** | **Haploids** | **HIR (%)** |
| --- | --- | --- | --- | --- | --- |
| DF199 | AC34-T2-2 | 26.08 | 1,536 | 19 | 1.24 |
|  | AC34-T2-4 | 24.62 | 927 | 14 | 1.51 |
|  | AC34-T2-15 | 26.31 | 751 | 14 | 1.86 |
|  | AC34-T2-16 | 26.12 | 1,084 | 13 | 1.20 |
| MT.AC | AC34-T2-4 | 15.56 | 1,688 | 24 | 1.42 |
|  | AC34-T2-9 | 24.69 | 1,483 | 21 | 1.42 |
|  | AC34-T2-10 | 21.81 | 677 | 9 | 1.33 |
|  | AC34-T2-15 | 19.33 | 2,430 | 39 | 1.60 |
|  | AC34-T2-16 | 16.53 | 1,880 | 32 | 1.70 |
| DF1.MT | AC34-T2-4 | 18.73 | 1,358 | 34 | 2.50 |
|  | AC34-T2-13 | 18.65 | 1,982 | 38 | 1.92 |
|  | AC34-T2-15 | 22.44 | 749 | 24 | 3.20 |
| JZ801 | AC34-T2-13 | 15.67 | 967 | 19 | 1.96 |
|  | AC34-T2-41 | 15.22 | 379 | 5 | 1.32 |
| 1004 | AC34-T2-2 | 21.27 | 148 | 4 | 2.70 |
|  | AC34-T2-15 | 28.51 | 246 | 4 | 1.63 |
| 1006 | AC34-T2-15 | 26.00 | 335 | 9 | 2.69 |
|  | AC34-T2-16 | 23.13 | 344 | 8 | 2.33 |
| 4013 | AC34-T2-Mix | 28.74 | 693 | 6 | 0.87 |
| 4014 | AC34-T2-Mix | 12.57 | 357 | 2 | 0.56 |
| 4015 | AC34-T2-Mix | 9.97 | 809 | 4 | 0.49 |
| 4024 | AC34-T2-Mix | 24.50 | 191 | 2 | 1.05 |
| 4025 | AC34-T2-Mix | 34.12 | 962 | 5 | 0.52 |
| 4029 | AC34-T2-Mix | 22.12 | 252 | 2 | 0.79 |
| 4034 | AC34-T2-Mix | 18.48 | 437 | 7 | 1.60 |
| 4035 | AC34-T2-Mix | ND | 307 | 8 | 2.61 |
| 4036 | AC34-T2-Mix | ND | 226 | 7 | 3.10 |
| 4037 | AC34-T2-Mix | ND | 148 | 2 | 1.35 |
| 4038 | AC34-T2-Mix | ND | 137 | 4 | 2.92 |
| 4039 | AC34-T2-Mix | ND | 151 | 4 | 2.65 |
| 4040 | AC34-T2-Mix | ND | 190 | 7 | 3.68 |
| 4041 | AC34-T2-Mix | ND | 201 | 3 | 1.49 |
| 4042 | AC34-T2-Mix | ND | 332 | 8 | 2.41 |
| 4043 | AC34-T2-Mix | ND | 485 | 9 | 1.86 |
| 4044 | AC34-T2-Mix | ND | 360 | 9 | 2.50 |
| 4048 | AC34-T2-Mix | ND | 75 | 1 | 1.33 |
| 4049 | AC34-T2-Mix | ND | 440 | 7 | 1.59 |
| 4050 | AC34-T2-Mix | ND | 331 | 5 | 1.51 |
| 4053 | AC34-T2-Mix | ND | 454 | 7 | 1.54 |
| 4055 | AC34-T2-Mix | ND | 182 | 5 | 2.75 |
| 4056 | AC34-T2-Mix | ND | 268 | 7 | 2.61 |
| 4057 | AC34-T2-Mix | ND | 369 | 9 | 2.44 |
| 4059 | AC34-T2-Mix | ND | 238 | 7 | 2.94 |
| 4060 | AC34-T2-Mix | ND | 233 | 6 | 2.58 |
| 4063 | AC34-T2-Mix | ND | 522 | 14 | 2.68 |
| 4064 | AC34-T2-Mix | ND | 576 | 11 | 1.91 |
| 4066 | AC34-T2-Mix | ND | 354 | 9 | 2.54 |
| 4068 | AC34-T2-Mix | ND | 153 | 2 | 1.31 |
| **Total** | **AC34-T2** | **24.70** | **29,397** | **509** | **1.94 ± 0.74** |

Haploid plants are sterile and must undergo chromosome doubling to develop into fertile plants. Chromosome doubling is usually induced chemically, but spontaneous doubling of haploid embryos *in vitro* is also commonly observed (Seguĺ-Simarro and Nuez, 2008). Surprisingly, spontaneous diploidization was also observed *in vivo* in *dmp*-induced tomato haploid plants, as evidenced by the production of viable pollen and seed-bearing fruits (Supplemental Figure 10). All nine haploid plants produced viable pollen, of which seven (78%) produced fruits and three (33%) eventually produced viable seeds (Supplemental Table 8). Tomato haploid plants can be easily propagated using cuttings from the parent plant (Supplemental Figure 11) and might be used to generate a higher proportion of spontaneous DH plants. After treating cuttings with the chemical doubling agent colchicine, three out of seven haploid plants successfully converted into DH plants with diploid cells and a high percentage (62%) of viable pollen (Supplemental Figure 12). Overall, we established a tomato DH breeding method including *DMP-*HI, FAST-Red based haploid identification and chromosome doubling.

### *DMP*-HI systems for dicot polyploid crops

To determine whether the *DMP-*HI system could be used in polyploid dicots crops, we evaluated the system in rapeseed (cv. Westar) and tobacco (cv. K326), both of which are amphidiploids. We found that one of the possible four *BnDMP* genes, *BnDMP1C,* was lost from the Westar genome, which was confirmed by a BLAST search (Song et al., 2020). Knock-out mutants of the three *DMP* genes in rapeseed (Figure 4A and 4B and Supplemental Table 3) and tobacco were obtained using CRISPR-Cas9 mediated mutagenesis (Figure 5A and 5B and Supplemental Table 3). We first determined the seed setting rate in *B. napus* selfing and crossing progenies. Compared with wild type, the number of filled seeds and the percentage seed set was significantly reduced in *bndmp* triple mutants (Figure 4C and 4D). Seed setting rate was not evaluated in tobacco.

**Figure 4.**
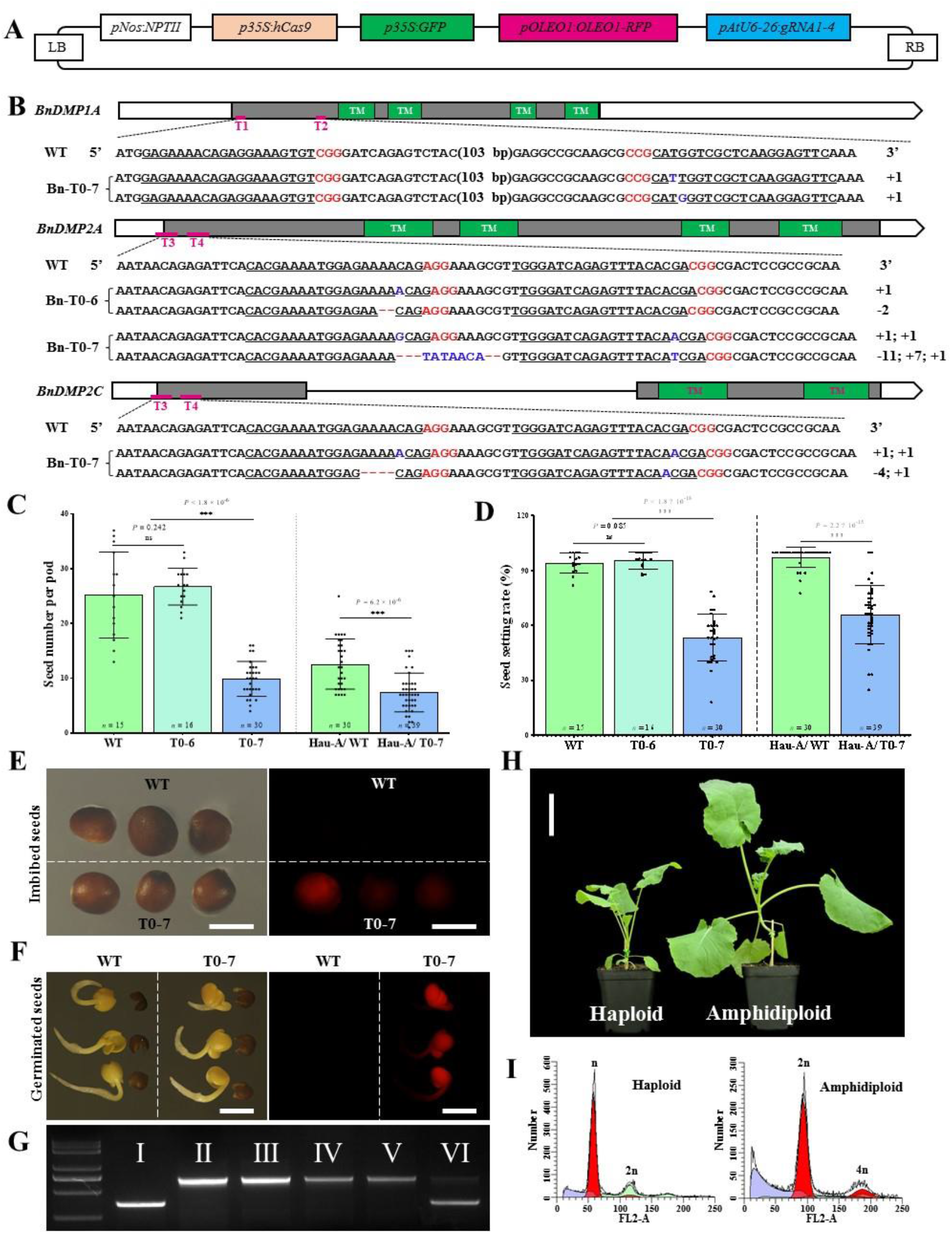
Mutation of BnDMP genes induces haploid seed formation. (**A**) The CRISPR/Cas9 mutagenesis vector comprising four sgRNAs (*gRNA1-4*) targeting three *BnDMP* genes, and the *pNos:NPTII* and the *p35S:GFP* and FAST-Red (*pOLEO1:OLEO1-RFP*) selection cassettes. (**B**) Schematic representation of the wild-type *BnDMP* genes. Filled blocks, clear blocks and the gray line indicate the coding region, the untranslated regions, and the intron, respectively. Green blocks correspond to the four predicted transmembrane domains (TM). Pink lines indicate the regions (T1, T2, T3 and T4) targeted by the sgRNAs. The sequences from wild type (WT) and mutant alleles are shown below the overview. The sgRNA target sequences are underlined, and the protospacer-adjacent motif (PAM) is shown in red. Nucleotide insertions are shown in blue and deletions by red dashes. (**C** and **D**) Quantification of seed number per pod (C) and seed set phenotypes (D) from WT and independent CRISPR–Cas9 lines. Data represent the mean ±s.d.; \*\*\**p* < 0.001 (two-tailed Student's *t*-test); ns, not statistically significant. *n*, number of siliques. (**E** and **F**) RFP expression of WT and T0-7 transgenic seeds in the imbibed (E) and germinated (F) state under white light (left panel) and fluorescent light (right panel). (**G**) Seedlings lacking RFP expression in germinated embryos were genotyped with polymorphic markers between the inducer line and testers. The left lane shows the DNA size marker, and the Roman numbers I to VI represent the PCR products from the inducer (I), Hau-A (II), haploid (III to V), and amphidiploid (VI) lines. (**H**) Representative images of rapeseed haploid and amphidiploid seedlings. (**I**) Flow cytometry verification of the ploidy of a putative haploid and an amphidiploid control. The *x* axis represents the signal peak for the nucleus and the *y* axis represents the number of nuclei. Scale bars: 5 mm (E and F) and 5 cm (H). In E to I, experiments were repeated at least three times and similar results were obtained.

**Figure 5.**
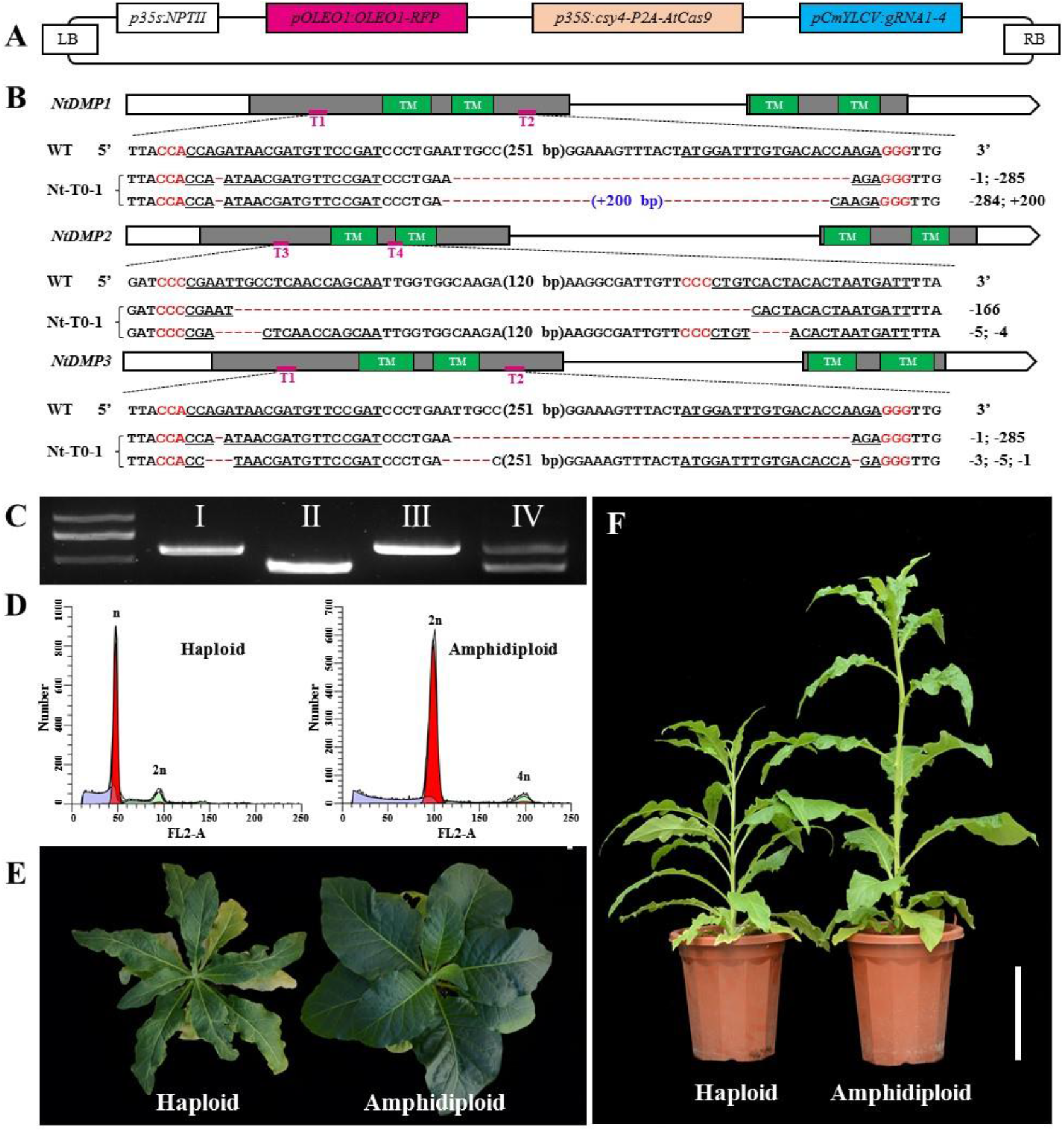
Tobacco *ntdmp* mutation induces maternal haploids. (**A**) Schematic of the *NtDMP* CRISPR/Cas9 mutagenesis vector. (**B**) Schematic representation of the wild-type (WT) *NtDMP* genes. Filled blocks, clear blocks and the gray line indicate the coding region, the untranslated regions, and the intron, respectively. Green blocks correspond to the four predicted transmembrane domains (TM). Pink lines indicate the regions (T1, T2, T3 and T4) targeted by the sgRNAs. The sequences from WT and mutant alleles from the three tobacco *DMP* genes are shown below the overview. The sgRNA target sequences are underlined, and the protospacer-adjacent motif (PAM) is shown in red. Nucleotide insertions are shown in blue and deletions by red dashes. (**C**) Seedlings from cross progeny were genotyped with polymorphic markers between the inducer line and testers to identify putative haploids. The left lane shows the DNA size marker, and the Roman numbers I to IV represent the PCR products from the K326 (I), inducer (II), haploid (III) and amphidiploid (IV) lines. (**D**) Flow cytometry verification of the ploidy of a putative haploid and an amphidiploid control. The *x* axis represents the signal peak for the nucleus and the *y* axis represents the number of nuclei. (**E** and **F**) Representative images of tobacco haploid and tetraploid seedling (E) and plants (F). Scale bars: 10 cm (E) and 20 cm (F). In E to F, experiments were repeated at least three times and similar results were obtained.

Next, we further verified the HI ability of the *bndmp* and *ntdmp* mutants. First, selfed progenies of *bndmp* and *ntdmp* mutants were screened for putative haploids based on their phenotypes. One of 97 T_1_ *bndmp* triple mutant plants (1.0%) and nine plants of 1,111 T_1_ *ntdmp* triple mutant plants (0.8%) showed the typical haploid phenotype (Table 2 and Table 3 and Supplemental Figure 13 and Supplemental Figure 14). These putative haploid plants were subsequently confirmed to be true haploids by ploidy analysis, suggesting that *bndmp* and *ntdmp* loss-off-function mutant can induce haploids in selfed progenies.

**Table 2.**
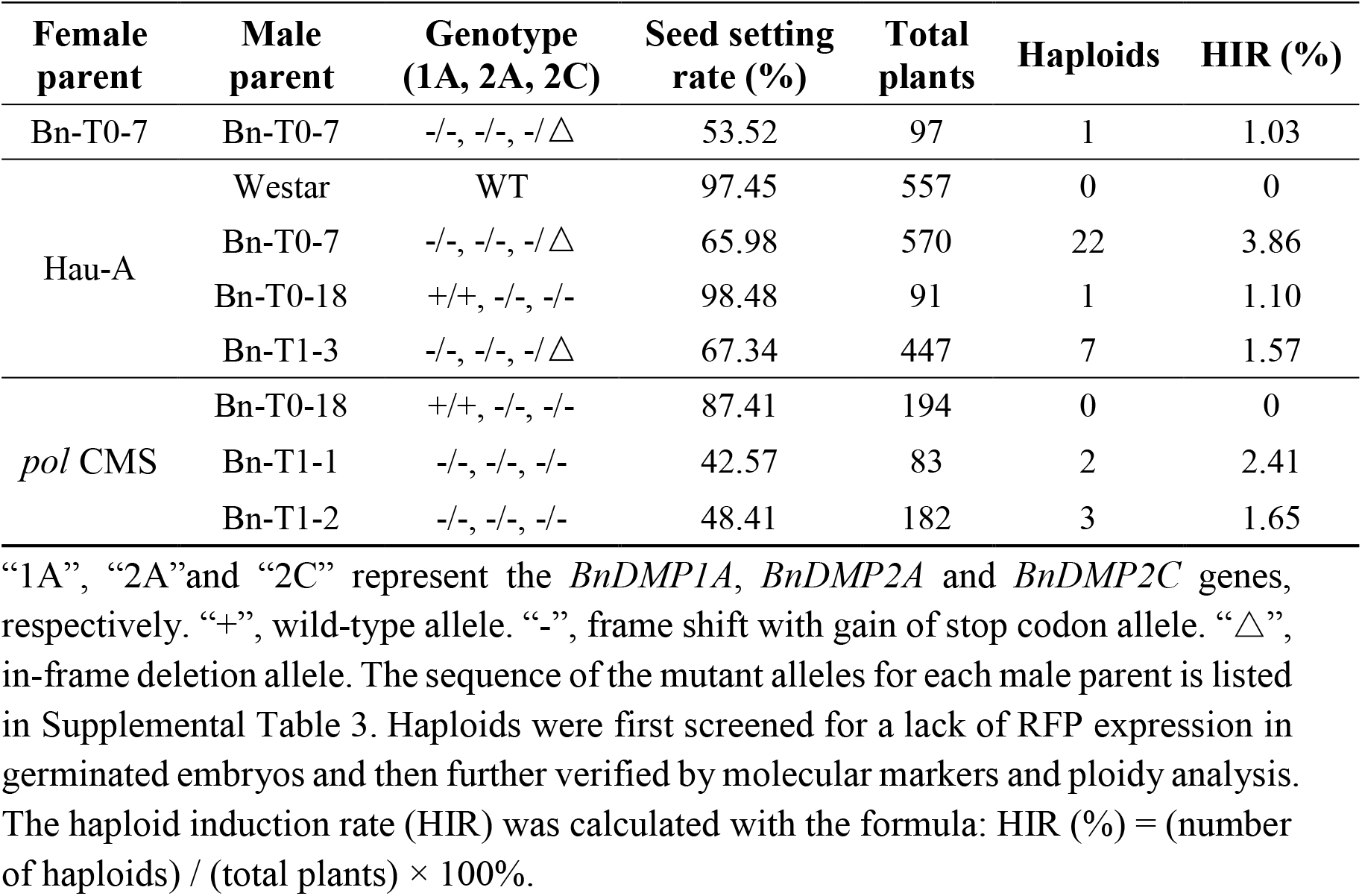
Haploid induction rate in *bndmp* mutant progeny.

**Table 3.**
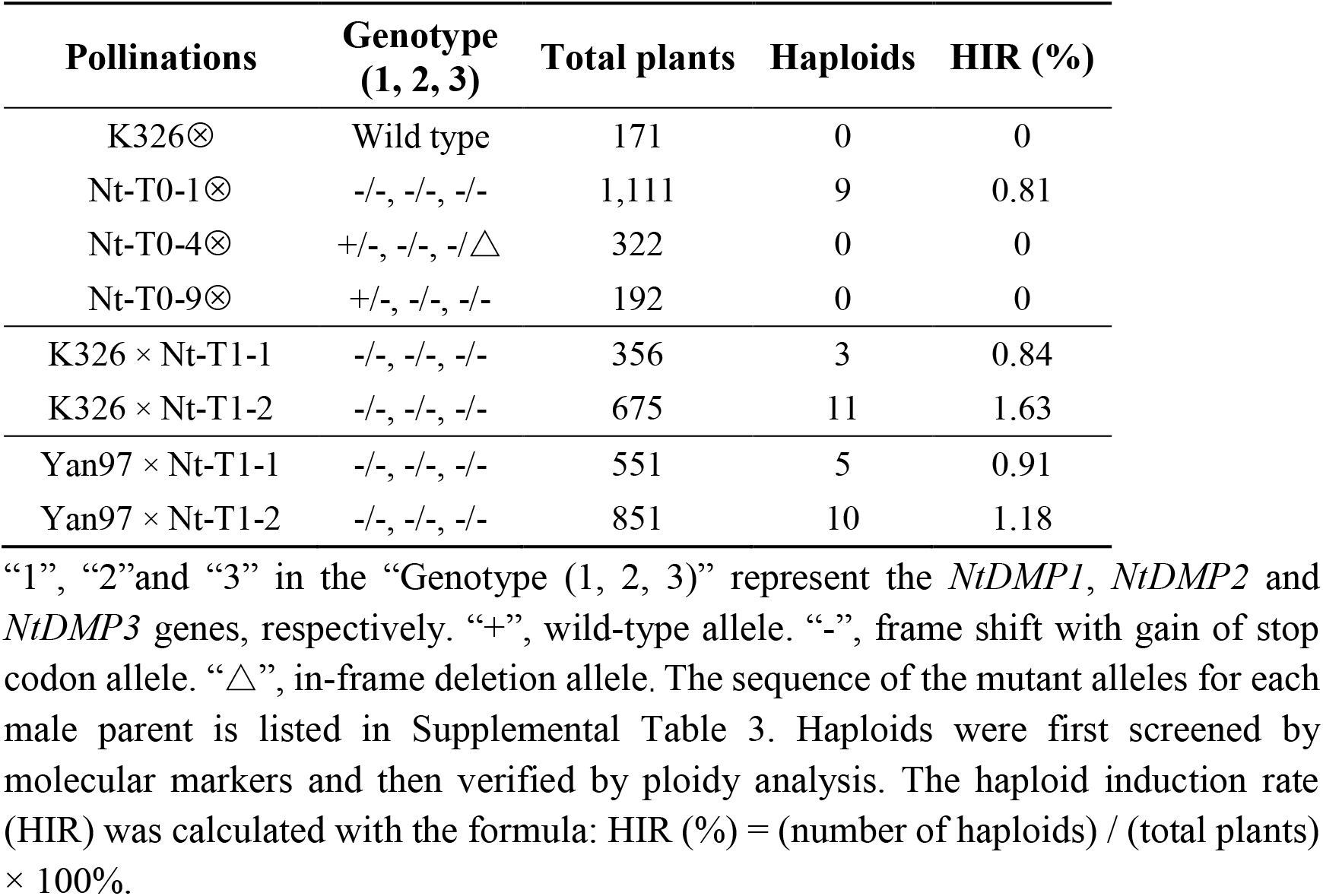
Haploid induction rate in *ntdmp* mutant progeny.

Then, crosses between the cytoplasmic male sterile lines Hau-A and *pol* CMS with *bndmp* mutants and between cultivars K326 and Yan97 with *ntdmp* mutants were made to determine whether *dmp* mutations can induce maternal haploids when used as the male parent. Unlike tomato, both rapeseed and tobacco only showed weak FAST-Red expression in imbibed seeds (Figure 4E), which made it difficult to identify haploid embryos at this stage. Therefore, all crossed progenies were germinated in water and the embryos assayed for RFP expression. The FAST-Red marker was not expressed in germinating embryos from the *ntdmp* mutant crosses (data not shown), but was expressed in cotyledons of germinated embryos from the *bndmp* mutant crosses (Figure 4F). Therefore, rapeseed putative haploids were first screened with Fast-Red marker and then further evaluated by molecular marker analysis (Figure 4G), while tobacco putative haploids were screened with molecular markers (Figure 5C). Putative haploid plants were subsequently confirmed by ploidy analysis and plant phenotype (Figure 4, H and I and Figure 5D-5F and Supplemental Figure 13 and Supplemental Figure 14). Overall, we found that crossing with *dmp* mutants induces a HIR of 1.1% to 3.9% in *B. napus* (Table 2) and a HIR of 0.8% to 1.6% in tobacco (Table 3).

## DISCUSSION

Our study demonstrates for the first time that *dmp* mutants induce *in vivo* maternal haploids in multiple dicot crops. More importantly, we also show that *dmp* mutation can be used for haploid induction in a wide range of genotypes: *dmp* mutations in three tomato genotypes were used to obtain haploids in 39 different female genotypes that differ in their genetic backgrounds, including both determinate and indeterminate growth types, as well as fruit types that differ in color, shape and size (Supplemental Table 4). Our data therefore suggest that a single *dmp* mutant line can be used to develop a genotype-independent DH technology in dicot crops.

Identification of the correct ZmDMP (co)ortholog for HI can be challenging due to low sequence identity and/or the presence of multigene families. In this study, we show that candidate *DMP* genes can be initially selected based on their sequence identity and by their pollen/flower expression pattern, and then verified in a complementation strategy in arabidopsis. Although we did not assess *DMP*-HI in chili pepper, cotton, soybean and cucumber, our success in developing a *DMP*-HI system in tomato, rapeseed and tobacco suggests that this bioinformatics approach can be used to identify candidate *DMP* genes for the development of HI systems in other dicots.

Given the presence of conserved *DMP* genes in dicot species (Zhong et al., 2020), and our success in inducing haploids in tomato, rapeseed and tobacco, it is likely that *DMP* mutation can be applied easily to generate *in vivo* haploid inducers in other dicot crops. Most commercial varieties of self-compatible vegetable crops are F_1_ hybrids. For many crops, up to 100% of a professional company’s seed portfolio can comprise F_1_ hybrids. F_1_ hybrid production requires the development of near homozygous parent lines, which can be greatly accelerated using DH technology, but often no widely-applicable *in vitro* or *in vivo* DH protocols are available, as is the case for tomato (Jacquier et al., 2020; Hooghvorst and Nogués, 2020a). In addition to self-compatible crops, the *DMP-*HI system might useful to induce maternal haploids via outcrossing for self-incompatible species like tetraploid potato (Ye et al., 2018), for which it is difficult to produce inbred lines for breeding. Moreover, *DMP* genes can also be found in fruit and forest trees species, like apple (*Malus domestica*), sweet cherry (*Prunus avium*) and rubber tree (*Hevea brasiliensis*), which take many years to reach the reproductive stage and for which homozygous line production is a lengthy process. *DMP*-HI systems can also be developed in non-transformable crops by using chemical/radiation mutagenesis to generate *dmp* mutants. Development of a genotype-independent *DMP*-HI system to other dicot crops would therefore represent a major advance over *in vitro* haploid production, where individual protocols must be developed for every species and genotype. This is especially true for members of the Solanaceae, Fabaceae, and Cucurbitaceae where recalcitrance for DH production is a major bottleneck for efficient breeding (Hooghvorst and Nogués, 2020a).

The FAST-Red marker facilitated efficient identification of maternal haploids in tomato. However, other than in arabidopsis (Zhong et al., 2020) and tomato, FAST-Red was not observed in rapeseed seeds, which might be due to the thick and darker seed coat of rapeseed, but could be observed in germinating embryos. Given that the same marker was only weakly expressed in tobacco seeds, either a more highly expressed late seed promoter or other visible markers, e.g., RUBY system (He et al., 2020), could be evaluated in future study to set robust haploid identification system. Alternatively, haploids can be selected based on the absence of paternal morphological or molecular markers.

We observed that the HIR in tomato is influenced by the female genotype (Table 1), as reported previously in maize (Prigge et al., 2011; Wu et al., 2014). This implies that beside novel paternal enhancers, novel maternal enhancers can also be identified and used in combination with *DMP* mutation to develop even more robust *in vivo* HI protocols. Given the universality of the *DMP*-HI system, enhancer screens for improved HIR could be carried out in any one crop, and newly identified enhancer genes implemented in other crops.

## METHODS

### Identify *DMP* genes in dicot crops

The full-length amino acid sequence of ZmDMP was first used for a BLASTP search (https://blast.ncbi.nlm.nih.gov/Blast.cgi) to identify the *DMP* gene with the highest identity for each crop. Then, these *DMP* genes were used as query to search the corresponding genome database of each crop and the *DMP* genes with >50% amino acid identity were selected for further analysis. Domain and transmembrane helices prediction was performed using the Pfam database (http://pfam.xfam.org) and TMHMM - 2.0 (https://services.healthtech.dtu.dk/service.php?TMHMM-2.0), respectively. Expression patterns of *DMP* genes were obtained from published data (Qin et al., 2014; Zhong et al., 2020) and public databases (https://biodb.swu.edu.cn/brassica/home and http://cucurbitgenomics.org/). The full-length amino acid sequences of *DMP* genes were aligned with MUSCLE embedded in SnapGene software (from Insightful Science; available at snapgene.com). Detailed information on the *DMP* genes is provided in Supplemental Table 1.

### Vector construction

Primers used to amplify and sequence the *DMP* alleles were designed based on the *DMP* gene sequences downloaded from Gramene or NCBI. For rapeseed and tomato, all gRNAs were driven by the *U6-26* promoter. *NPTII* driven by the *nopaline synthase* (*Nos*) promoter from *Agrobacterium tumefaciens* and *35S:GFP* were used for transgene selection during transformation, and, FAST-Red was used for identification of haploid seeds. These cassettes, together with human codon-optimized Cas9 driven by *35S* promoter, were introduced simultaneously in one step into pISCL4723 through the Golden Gate cloning method (Wang et al., 2019). To generate the CRISPR/Cas9 mutagenesis construct for tobacco *DMP* genes, the FAST-Red cassette was first amplified with FAST-Red-F/R primers and ligated into KpnI-linearized pDIRECT_22C to yield pDIRECT_22C_FastR using the Seamless Assembly Cloning Kit (C5891–25, Clone Smarter). Four gRNAs separated by 20-bp Csy4-binding sites were introduced simultaneously in one step into pDIRECT_22C_FastR with a previously reported protocol (Čermák et al., 2017). For *DMP* gene complementation constructs, CDS sequences of each *DMP* gene from different crops were cloned and driven by the *AtDMP9* promoter (1836 bp upstream of the ATG start codon). FAST-Red and *35S:RUBY* (He et al., 2020) were used for transgenic seed selection. These cassettes were introduced simultaneously in one step into pISCL4723 through the Golden Gate cloning. The primers used for vector construction are listed in Supplemental Table 9.

### Plant materials

The spring-type rapeseed cultivar Westar, a tobacco cultivar K326 and three tomato cultivars (Ailsa Craig, Micro-Tom and Moneyberg) were used as receptor lines to knock out *DMP* genes. Two male sterile lines of rapeseed (Hau-A and *pol* CMS), two cultivars of tobacco (K326 and Yan97) and dozens of tomato varieties (described in Supplemental Table 4) were used as the female parents to evaluate the outcrossing HIR of *dmp* mutants. Rapeseed, tobacco and tomato transgenic plants were obtained by *Agrobacterium-mediated* transformation as previously described (Horsch et al., 1985; van Roekel et al., 1993; Dai et al., 2020). All plants were grown in the greenhouse under natural light. The arabidopsis *dmp8dmp9* mutant (Zhong et al., 2020) was used to generated different complementation lines by *Agrobacterium tumefaciens*-mediated floral dip transformation (Clough and Bent, 1998).

### Screening for *dmp* mutants

Sanger sequencing was performed to identify *dmp* mutants in T0 transgenic lines. All sequences were aligned to the *DMP* wild-type allele with SnapGene software. PCR products containing multiple amplification products from the same locus were further amplified with KOD FX (Toyobo) and cloned into the *pEASY* vector (pEASY-Blunt Zero Cloning Kit, TransGen Biotech). At least six independent colonies were selected and sequenced by the Tsingke Biological Technology Co., Ltd. The primers used for *dmp* mutants genotyping are list in Supplemental Table 9.

### Pollen viability evaluation

Mature tomato flowers were used for the pollen analyses. A needle was used to slice open the anther lengthwise and then dragged upwards through the locule of the anther to collect pollen on the tip of the needle. For the pollen germination experiment, pollen grains were incubated in liquid medium (10% sucrose, 0.01% boric acid, 0.1% yeast extract, 5 mM CaCl_2_, 50 μM KH_2_PO_4_ and 15% PEG 4000) in the dark at 30 °C for 1 hour. Images were captured using a light microscope (CX41, Olympus, Japan) fitted with a Nikon DS-Ri1 camera. Viability was scored using the pollen germination rate. A pollen grain was scored as germinated when it formed a pollen tube. For the pollen staining experiment, pollen grains were stained on a microscope slide using Alexander’s stain solution (Solarbio) and photographed using a light microscope (Axio Imager Z2, Zeiss, Germany) fitted with a Canon EOS 6D camera. Red-stained pollen was scored as viable.

### Analysis of seed phenotypes

Tomato and rapeseed seeds were harvested from ripe fruits and used to score phenotypes. Seeds were divided into three categories (normal seeds, aborted seeds and undeveloped ovules) based on their size and color. For tomato seed dissection, a razor blade was used to cut the imbibed seeds lengthwise into two uneven pieces (1/4 and 3/4). The larger piece containing the embryo was used to separate the embryo and testa from the endosperm with fine forceps under a stereo microscope (S6D, Leica, Germany). Samples were observed and photographed using a stereo fluorescence microscope (SEX16, Olympus, Japan) fitted with Olympus DP72 camera. For arabidopsis, siliques were cleared with a previously described method (Zhong et al., 2020) and photographed using a Leica S6D stereo microscope.

### Haploid identification

Haploids from selfed *dmp* mutant lines were first identified based on their phenotype and then confirmed by flow cytometry.

In the T_1_ generation, *sldmp* mutant lines with or without the segregating Fast-Red marker were used as pollen donors in a cross with wild-type female parents. All the seeds derived from these crosses were sown in soil and grown to the seedling stage. One molecular marker with a polymorphism between the *sldmp* mutant lines and testers was used to screen for putative haploid seedlings. Then, the ploidy of these putative haploid seedlings was then confirmed by flow cytometry.

In the T_2_ generation, *sldmp* mutants with a homozygous Fast Red marker were used in crosses. Seeds derived from these crosses were first treated with 2% sodium hypochlorite for 15 minutes (to improve Fast-Red detection) and then divided into two groups (red seeds and white seeds) based on their color under white light. The red seeds with RFP expression in the embryo and endosperm were considered to carry diploid embryos. All the white seeds were treated a second time with 2% sodium hypochlorite for 15 minutes and sown on half-strength Murashige and Skoog medium. White seeds were further divided into two classes (strong RFP seeds and weak RFP seeds) under fluorescent light (excitation wavelength 540 nm, emission wavelength 600) using a hand-held lamp (LUYOR-3415RG). During germination, both seed classes were also checked for the absence/presence of RFP in the root tip. Seeds with weak RFP expression that did not show RFP signal in embryo root tip were scored as putative haploids. These putative haploids were further confirmed at the seedling stage by molecular markers and ploidy analysis.

In the wild-type × *bndmp* progeny, haploids were first screened with the Fast-Red marker in the cotyledon of germinated seeds under fluorescent light (excitation wavelength 540 nm, emission wavelength 600) using a hand-held lamp (LUYOR-3415RG) and molecular markers (A07-1), and then confirmed by flow cytometry.

In the wild-type × *ntdmp* progeny, both imbibed seeds and root tips of germinated seeds showed weak RFP expression, which made it difficult to identify haploids via Fast-Red marker. Therefore, all tobacco haploids were first screened by a molecular marker (*NtDMP2*) and then confirmed by flow cytometry. The primers used for identification of haploids are shown in Supplemental Table 9.

### Flow cytometry

Fresh leaves (0.5 g) from each sample were chopped with a razor blade in 2 mL lysis buffer as previously described (Zhong et al., 2020), and filtered through an 80 μm nylon filter. Nuclei were collected by centrifugation at 1000 r.p.m. for 5 minutes at 4 °C and stained with propidium iodide in the dark for 20 min. The ploidy level of each sample was analyzed with a BD FACSCalibur Flow Cytometer and BD CellQuest Pro software. Wild-type plants were used as a control and the position of its first signal peak was set at ~100 (FL2-A value). The samples with the first signal peak at ~50 (FL2-A value) were deemed to be haploids.

### Whole-genome resequencing and genotype calling of tomato haploids

Genomic DNA libraries of each sample were constructed and sequenced at an average depth of approximately 20-fold coverage using the Illumina high-throughput sequencing platform (Annoroad Gene Technology Co., Ltd, Beijing, China). Reads containing adapter sequence, reads with a high ratio of N (N bases accounting for more than 5% of the total reads) and reads with low quality (bases with a mass value less than 19 accounting for more than 50% of the total reads) were filtered from the raw data using fastp software(Chen et al., 2018). Clean reads of each sample were aligned to the tomato reference genome (SL4.0) using Bowtie2 software (Langmead and Salzberg, 2012). Uniquely mapped reads were used for SNP calling. Joint-genotype calling was carried out on the whole genome using HaplotypeCaller, CombineGVCFs and GenotypeGVCFs tools from GATK4 (ref.(Poplin et al., 2017)) (version 4.1.2.0). The SNPs with bi-alleles were selected and filtered with following parameters: QUAL < 1000.0, QD < 2.0, MQ < 40.0, FS > 60.0, SOR > 3.0, MQRankSum < −12.5, ReadPosRankSum < −8.0. These high quality SNPs were used for recombination map construction via a sliding window approach. The window size was set to 30 SNPs and the step size was set to 1 SNP.

### Chromosome dosage analysis of tomato haploids

The chromosome dosage analysis was performed as previously described (Tan et al., 2015). BAM files generated by Bowtie2 software were used to calculate the coverage of each bin from each sample with bamCoverage (parameters: --normalizeUsing CPM --binSize 100000) in deepTools (Ramírez et al., 2016) software. Relative coverage was calculated by dividing the coverage of each bin by the mean percentage of corresponding female parents.

### Propagation of tomato haploid plants from cuttings

Strong side shoots from haploid plants were cut from the parent plant with sharp scissors and placed in water containing 0.2% (*w*/*v*) rooting hormone powder (Shandong Huanuo Federal Agrochemical Co.,Ltd) for about two weeks under LED light (150 μmol m^-2^ s^-1^) on a 16 h light/8 h dark photoperiod. Cuttings with well-developed roots were transplanted into 6 L pots with soil and grown in the greenhouse under the natural light.

### Colchicine treatment of tomato haploids

For colchicine treatment, a solution was used that contained 0.06% colchicine (Cat#C3915, Sigma) dissolved in dimethyl sulfoxide, 2% DMSO, 5% glycerol and 1% Tween. For the control treatment, the same solution without colchicine was used. Haploid cuttings were planted and grown in the greenhouse under natural light for one month. The primary shoot apical meristem (SAM) was dipped in the colchicine solution for 1.5 min. The internode that was closest to the SAM was then marked as the starting point. After three to four new internodes developed from the treated meristem, expanded leaf samples were taken from the top internode for ploidy analysis.

## Data availability

The datasets generated and/or analysed during the current study are available from the corresponding author on request.

## FUNDING

This work was supported by National Key Research and Development Program of China (2016YFD0101200, 2018YFD0100201), China Agriculture Research System of MOF and MARA, National Natural Science Foundation of China (91935303, 32001554, 31991185), the 2020 Research Program of Sanya Yazhou Bay Science and Technology City (SKJC-2020-02-003), China Postdoctoral Science Foundation (2020TQ0356) and a China Scholarship Council PhD fellowship (201506350003).

## AUTHOR CONTRIBUTIONS

Y.Z., B.C., C.L., K.B. and S.C. conceived and designed the experiments. Y.Z., D.W., B.C., X.Z. and Y.W. performed most of the experiments. M.L., Y.L., J.Liu, J.Z., M.C., M.W., T.R., X.Q., D.C., Z.L., J.Li, C.C. and Y.J. performed some of the experiments. Y.Z., B.C., S.C., C.L., M.W. and W.L. analyzed the data. Y.Z., B.C., B.Y., S.H., K.B. and S.C. discussed and prepared the manuscript. All authors discussed the results and provided feedback on the manuscript.

## ACKNOWLEDGMENTS

We thank Prof. Wencai Yang and Huolin Shen for providing the tomato seeds, Prof. Pu Wang, Yongsheng Qiu, Qinnan Wang and Hailong Chang for providing the greenhouse, Dr. Yubing He for sharing the *RUBY* construct, Prof. Xiaohong Yang for help with the analysis of haploid sequence data, Zhongqiu Li for help with the amino acid alignment, and Yixiao Liu for drawing the pictures of tomato seeds and seedlings. We thank Charlotte Siemons, Mieke Weemen and Xiaobing Jiang for help with pollination experiments and Vera Veltkamp for sharing tomato FAST-Red information.

## Notes

### Competing Interest Statement

The authors have declared no competing interest.

