## Supplemental Figures and Tables for "A genotype independent *DMP*-HI system in dicot crops"

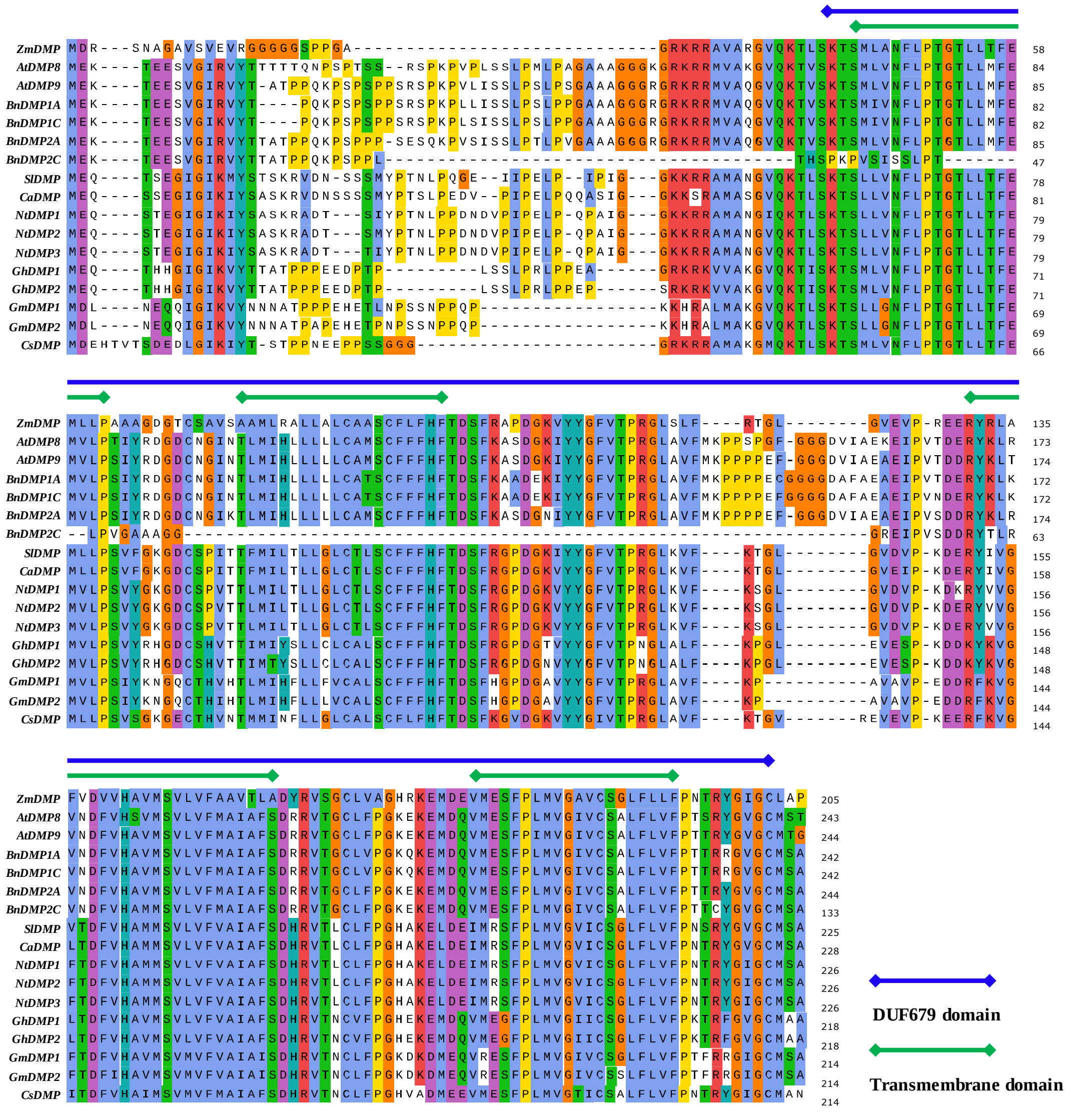


**Supplemental Figure 1. Amino acid alignment of DMP genes.** The alignment includes DMP proteins from maize (ZmDMP), arabidopsis (AtDMP8 and AtDMP9), Brassica napus (rapeseed) (BnDMP1A/BnaA03g55920D; BnDMP1C/BnaC03g03890D; BnDMP2A/BnaA04g09480D; BnDMP2C/BnaC04g31700D), tomato (SlDMP/Solyc05g007920), chili pepper (CaDMP/T459_27262), tobacco (NtDMP1/LOC107762412; NtDMP2/LOC107783066; NtDMP3/LOC107807404), cotton (GhDMP1/LOC107911807; GhDMP2/LOC107924398), soybean (GmDMP1/GLYMA_18G097400; GmDMP2/GLYMA_18G098300) and cucumber (CsDMP/Csa_1G267250). The DUF679 region is labeled by a blue line. Green lines indicate the four transmembrane domains predicted by TMHMM (https://services.healthtech.dtu.dk/service.php?TMHMM-2.0).

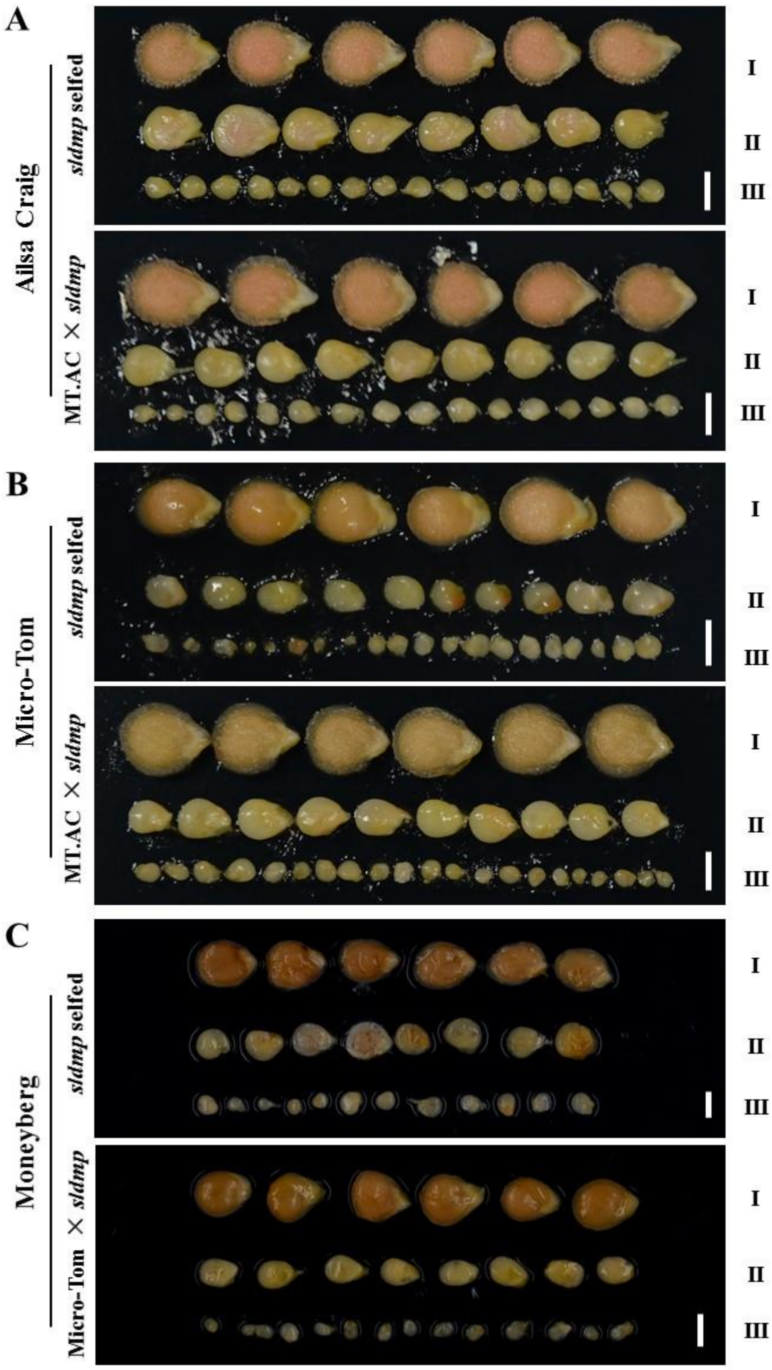


**Supplemental Figure 2. Seed phenotypes after *sldmp* selfing and outcrossing.** (**A** to **C**) Normal seeds (I), aborted seeds (II), and undeveloped ovules (III) after selfing and outcrossing *sldmp* mutants in the Ailsa Craig (**A**), Micro-Tom (**B**) and Moneyberg (**C**) backgrounds. Light images were taken of freshly harvested seeds from ripe fruits. MT.AC represents the $F_1$ hybrid of Micro-Tom × Ailsa Craig. Scale bar, 2 mm.

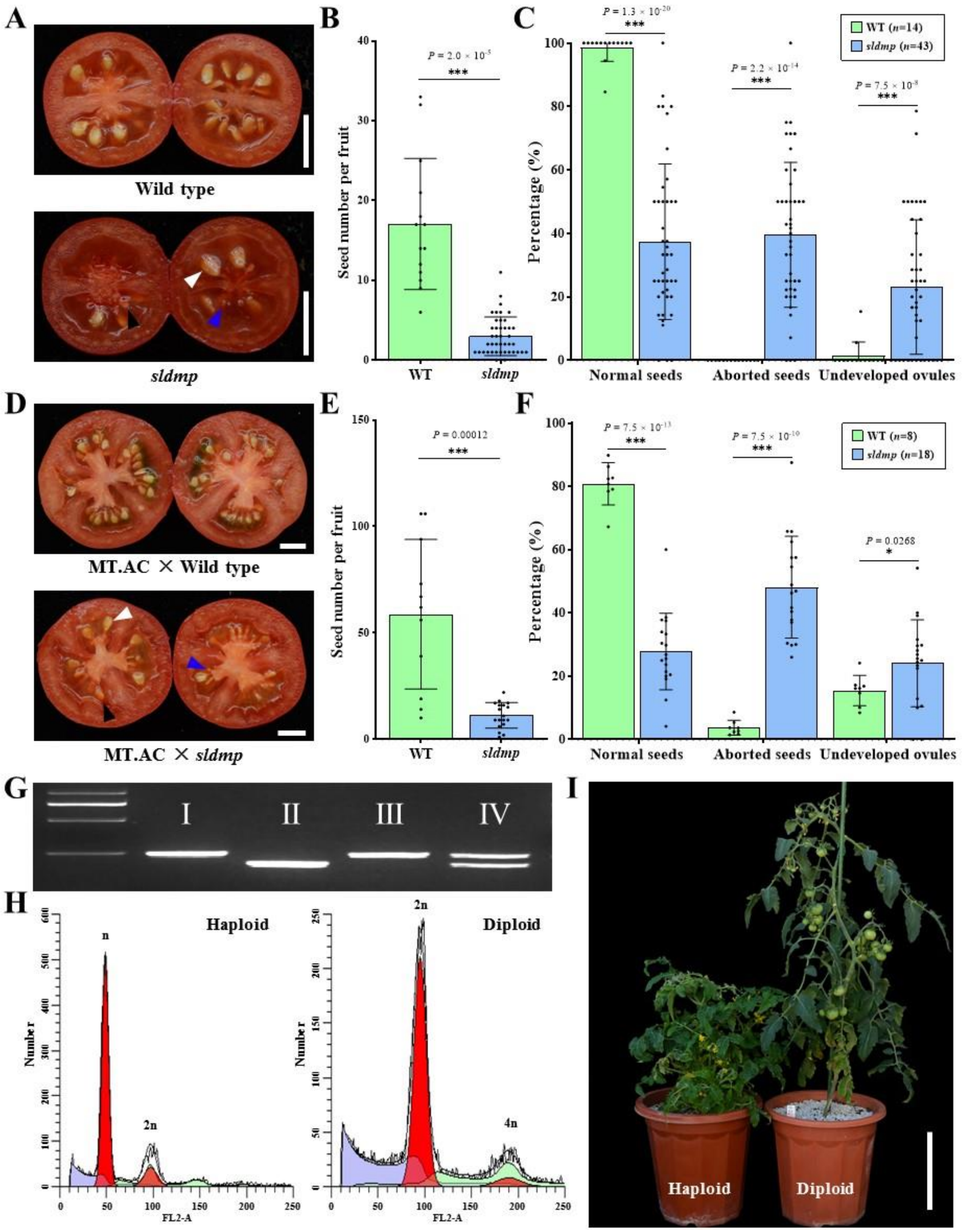


**Supplemental Figure 3. *SlDMP* mutation induces pleiotropic seed phenotypes in the Micro-Tom background.** (**A**) Representative images of ripened fruit from selfed wild type and a *dmp* mutant. White, black, and blue arrowheads indicate normal seeds, aborted seeds, and undeveloped ovules, respectively. (**B** and **C**) Quantification of seed number (B) and seed phenotypes (C) in fruits shown in (A). Data represent the mean ± s.d.; ****p* < 0.001 (two-tailed Student's *t*-test); *n*, number of fruits. d, Representative images of ripened fruit obtained after pollination by wild type and by the *sldmp* pollen. White, black, and blue arrowheads indicate normal seeds, aborted seeds, and undeveloped ovules, respectively. MT.AC, $F_1$ derived from Micro-Tom × Ailsa Craig. (**E** and **F**) Quantification of seed number (E) and seed phenotypes (F) in fruits shown in (D). Data represent the mean ± s.d.; **p* < 0.05, ****p* < 0.001 (two-tailed Student's *t*-test); *n*, number of fruits. (**G**) Identification of haploids and diploids using polymorphic markers for the Micro-Tom and Ailsa Craig backgrounds. The left-most lane shows the

DNA size marker. I, II, III and IV represent PCR products corresponding to Ailsa Craig, the *sldmp* mutant in the Micro-Tom background, haploid and diploid, respectively. (**H**) Flow cytometry verification of the ploidy of a putative haploid and a diploid control. The *x* axis represents the signal peak for the nucleus, and the *y* axis represents the number of nuclei. (**I**) Representative image of tomato haploid and diploid plant. Scale bars: 1 cm (A and D) and 20 cm (I).

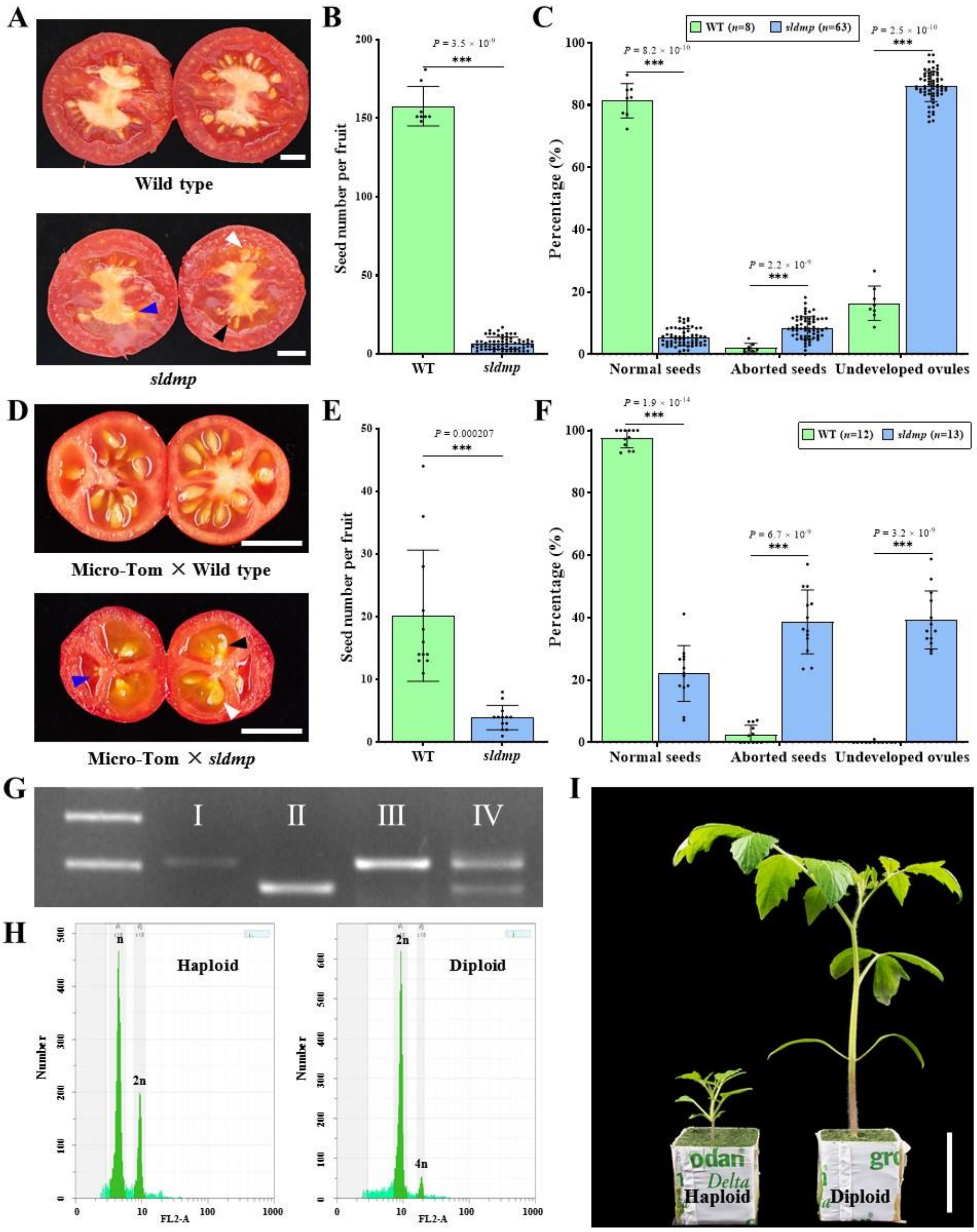


**Supplemental Figure 4.** ***SlDMP* mutation induces pleiotropic seed phenotypes in the Moneyberg background.** (**A**) Representative images of ripened fruit from selfed wild type and *dmp* mutant plants. White, black, and blue arrowheads indicate normal seeds, aborted seeds, and undeveloped ovules, respectively. (**B** and **C**) Quantification of seed number (B) and seed phenotypes (C) in fruits shown in (A). Data represent the mean ± s.d.; $***p < 0.001$ (two-tailed Student's *t*-test); *n*, number of fruits. (**D**) Representative images of Micro-Tom ripe fruits obtained after pollination by wild type and *sldmp* pollen. White, black, and blue arrowheads indicate normal seeds, aborted seeds, and undeveloped ovules, respectively. (**E** and **F**) Quantification of seed number (E) and seed phenotypes (F) in fruits shown in (D). Data represent the mean ±s.d.; $***p < 0.001$ (two-tailed Student's *t*-test); *n*, number of fruits. (**G**) Identification of haploids and diploids using polymorphic markers. The left-most lane shows the DNA size marker. I, II, III and IV represent PCR products corresponding to Micro-Tom, the *sldmp* mutant in the Moneyberg background, haploid and diploid, respectively. (**H**) Flow cytometry verification of the ploidy of a putative haploid and a diploid control. The *x* axis represents the signal peak for the nucleus, and the *y* axis represents the number of nuclei. (**I**) Representative image of tomato haploid and diploid seedlings. Scale bars: 1 cm (A and D) and 4 cm (I).

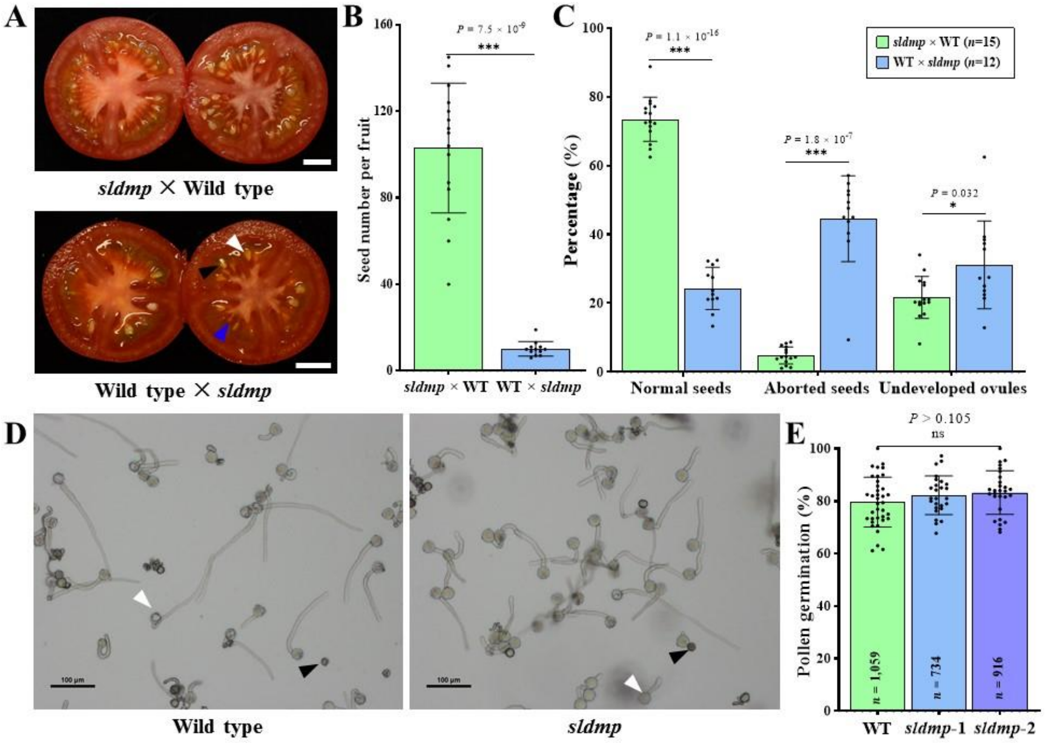


**Supplemental Figure 5. The paternal *sldmp* allele is responsible for *sldmp* mutant seed phenotypes.** (**A**) Representative images of ripened fruit from a reciprocal cross between *sldmp* and wild type, both in the Ailsa Craig background. White, black, and blue arrowheads indicate normal seeds, aborted seeds, and undeveloped ovules, respectively. (**B** and **C**) Quantification of seed number (B) and seed phenotypes (C) in fruits shown in (A). Data represent the mean ± s.d.; *$p < 0.05$, ***$p < 0.001$ (two-tailed Student's *t*-test); *n*, number of fruits. (**D**) Pollen viability of wild type and *sldmp* mutants in the Ailsa Craig background was determined by an *in vitro* pollen germination assay. Pollen with elongated tubes were scored as viable (white arrowhead) and pollen without tubes were scored as dead (black arrowhead). (**E**) Quantification of pollen viability, shown as the percentage of germinated pollen. Data represent the mean ± s.d.; ns, not statistically significant. *n*, the number of pollen grains. Scale bar, 1 cm (A) and 100 μm (D).

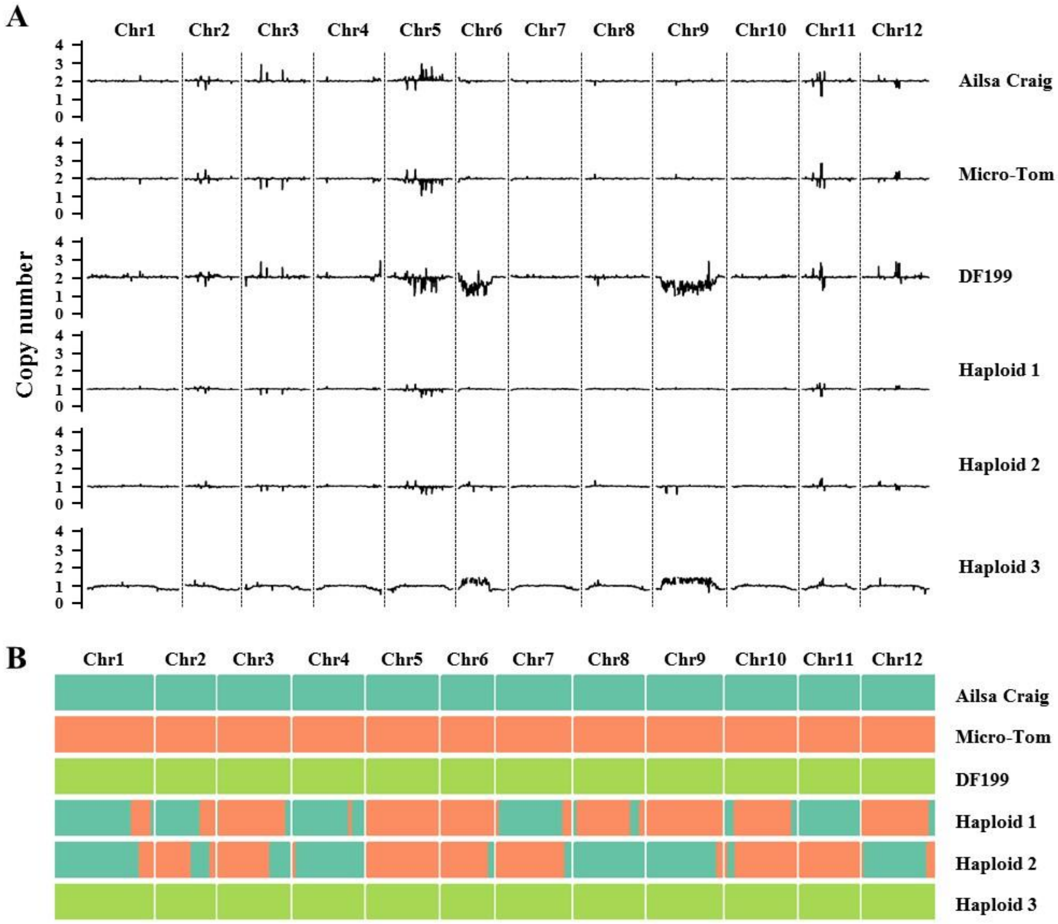


**Supplemental Figure 6. Dosage plots and genotype for tomato haploids.** (**A** and **B**) Dosage plots (A) based on 100 kbp non-overlapping bins and recombination maps (B) constructed via a sliding window approach across all twelve tomato chromosomes for three haploids and their corresponding parents. Sample names/IDs are indicated in the right of each plot. Haploids 1-3 were derived from MT.AC × *dmp* mutant in the Micro-Tom background, MT.AC × *dmp* mutant in the Ailsa Craig background and DF199 × *dmp* mutant in the Ailsa Craig background, respectively.

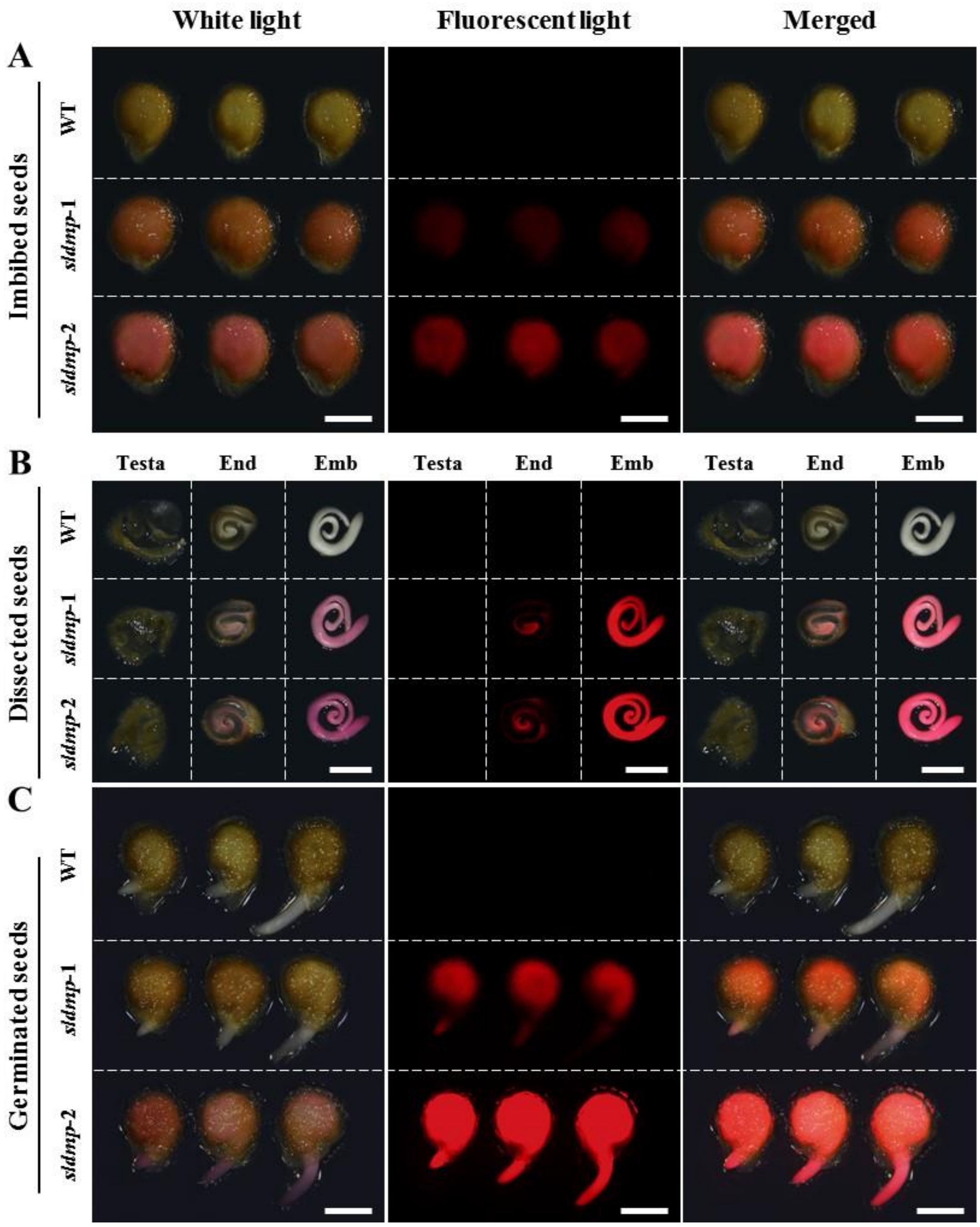


**Supplemental Figure 7. FAST-Red is expressed in the endosperm and embryo of tomato seeds.** (**A** to **C**) Analysis of RFP expression in the seeds from selfed wild type and two *dmp* mutants in the Ailsa Craig background. White light (left panel), florescent light (middle panel) and merged (right panel) micrographs of seeds in the imbibed (A), dissected (B) and germinated (C) state. End, endosperm. Emb, embryo. Scale bar, 2 mm. Experiments were repeated at least three times and similar results were obtained.

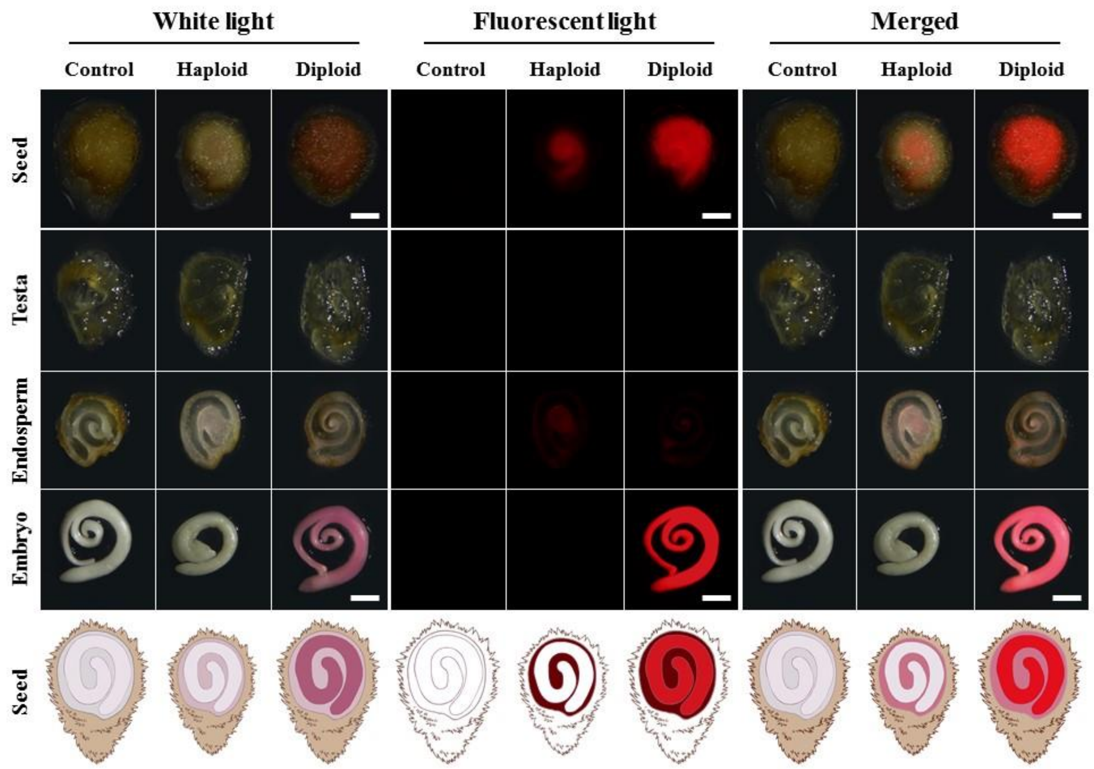


**Supplemental Figure 8. FAST-Red can be used as a visual marker to identify haploids in *sldmp* crosses.** The control seeds (DF199 × Ailsa Craig), haploid seeds (DF199 × *dmp* in the Ailsa Craig background, white seeds) and diploid seeds (DF199 × *dmp* in the Ailsa Craig background, red seeds) and their dissected contents (testa, endosperm and embryo) were observed under white light and fluorescent light. The bottom panel is a schematic representation of the different types of seeds shown in the panels directly above. Scale bar, 1 mm. Experiments were repeated at least three times and similar results were obtained.

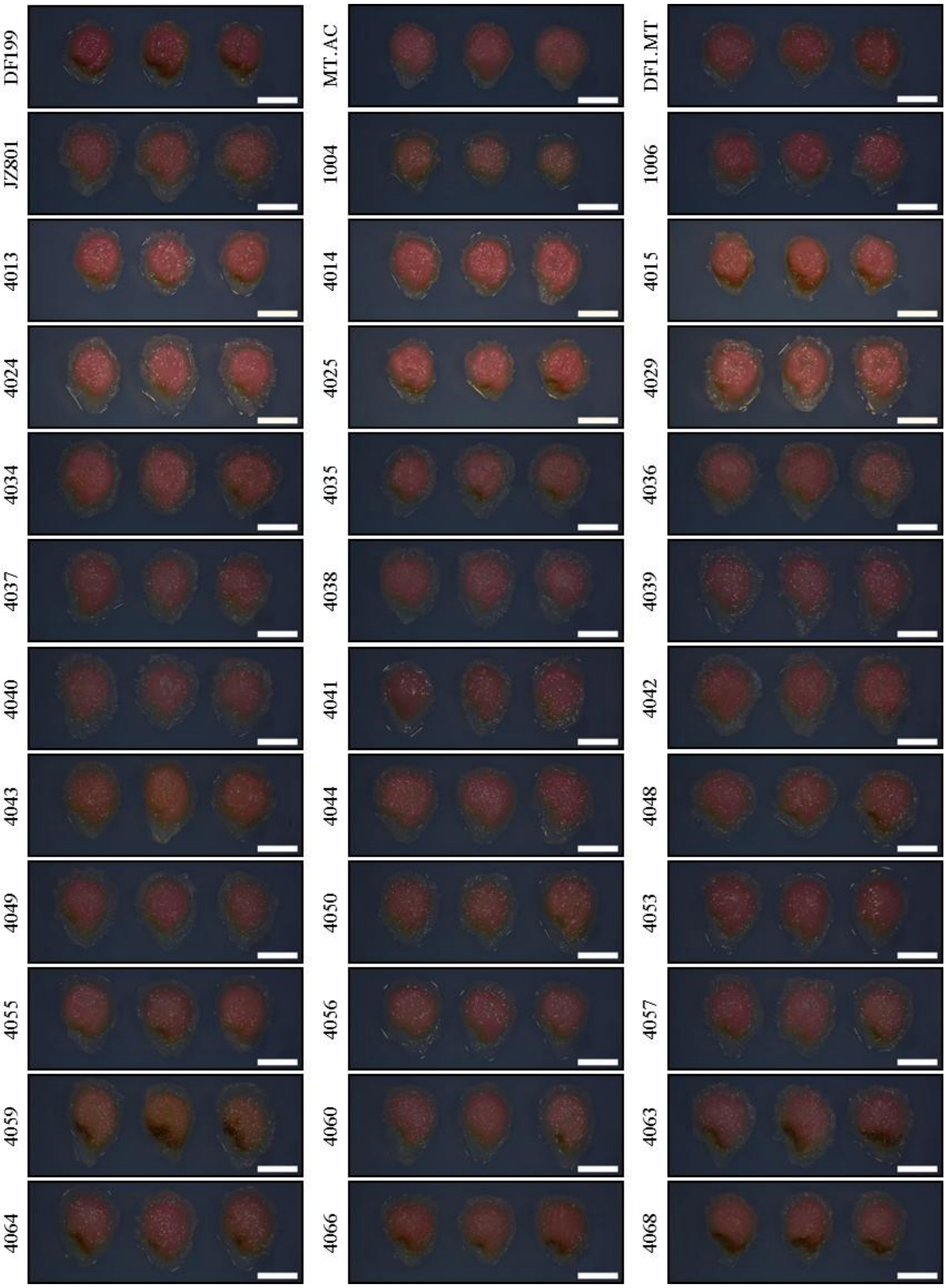


**Supplemental Figure 9. FAST-Red expression in different tomato backgrounds.** Representative images showing RFP expression in diploid seeds from 36 different hybrids pollinated by a *sldmp* mutant in the Ailsa Craig background. Hybrids names/IDs are indicated in the left side of each panel and detailed information of each hybrid is described in Supplemental Table 4. Scale bar, 2 mm. Experiments were repeated at least three times and similar results were obtained.

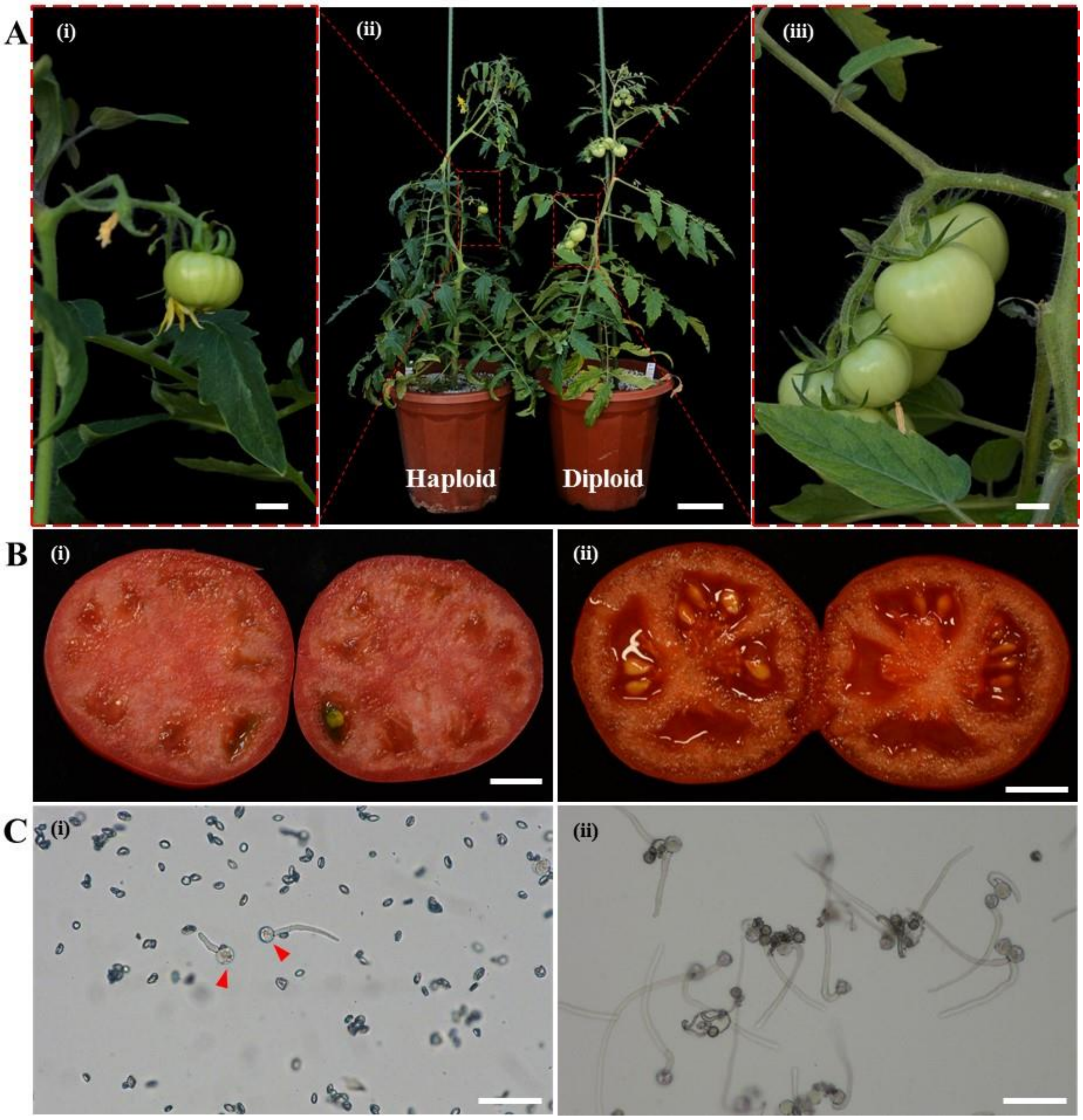


**Supplemental Figure 10. Spontaneous diploidization of haploid tomato plants.** (**A**) (ii) A representative image showing normal fruit development in a diploid plant and a haploid plant with sterile flowers and a fruit on the same inflorescence branch. The left and right panels are magnifications of the respective boxed regions, showing fruit formation in haploid (i) and diploid (iii) plants. (**B**) The ripened fruits from a haploid plant (i) with only a few seeds and from a diploid plant (ii) with many seeds. (**C**) Pollen viability in haploid (i) and diploid (ii) plants. The red arrowheads show that some pollen grains from the haploid plant can germinate, which could be due to spontaneous diploidization. Scale bar, 1 cm (A (i), A (iii) and B), 10 cm (A (ii)) and 100 μm (C).

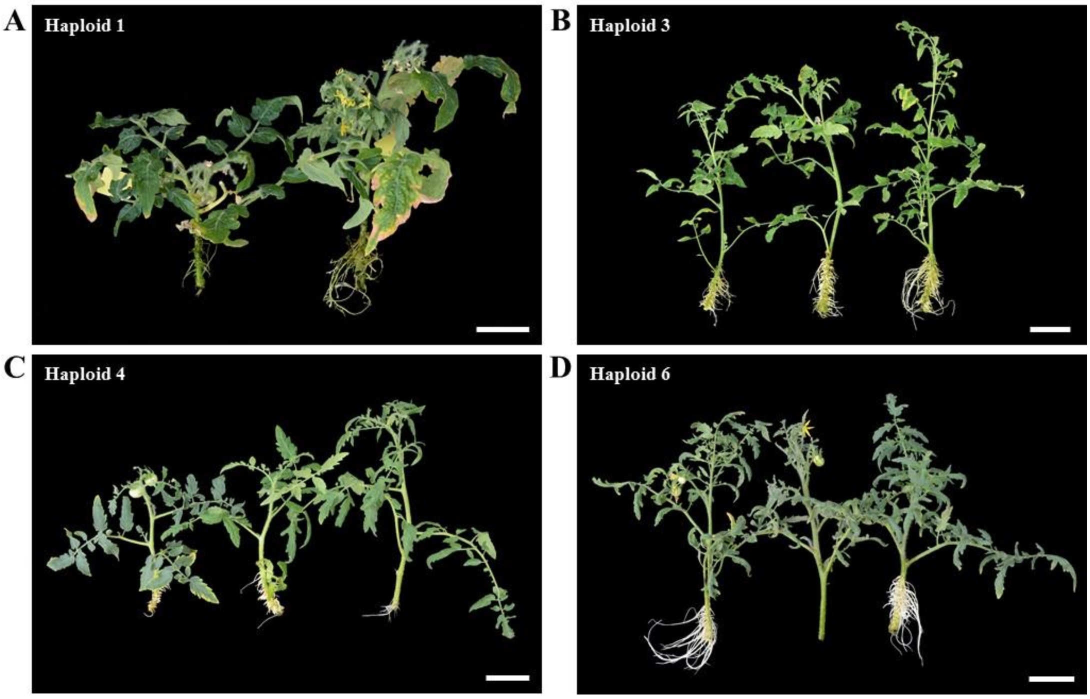


**Supplemental Figure 11. Haploid plant propagation by cuttings. (A** to **D**) Representative images of rooted cuttings from respectively *dmp*-induced haploid plants derived from the Micro-Tom × Ailsa Craig $F_1$ hybrid (A), DF199 (B), AF01 (C) and DF1 (D) backgrounds. Images are from cuttings that were placed for two weeks in rooting solution. The haploid plant identification codes are indicated in the top left corner of each images (Supplemental Table 8). Scale bar, 5 cm.

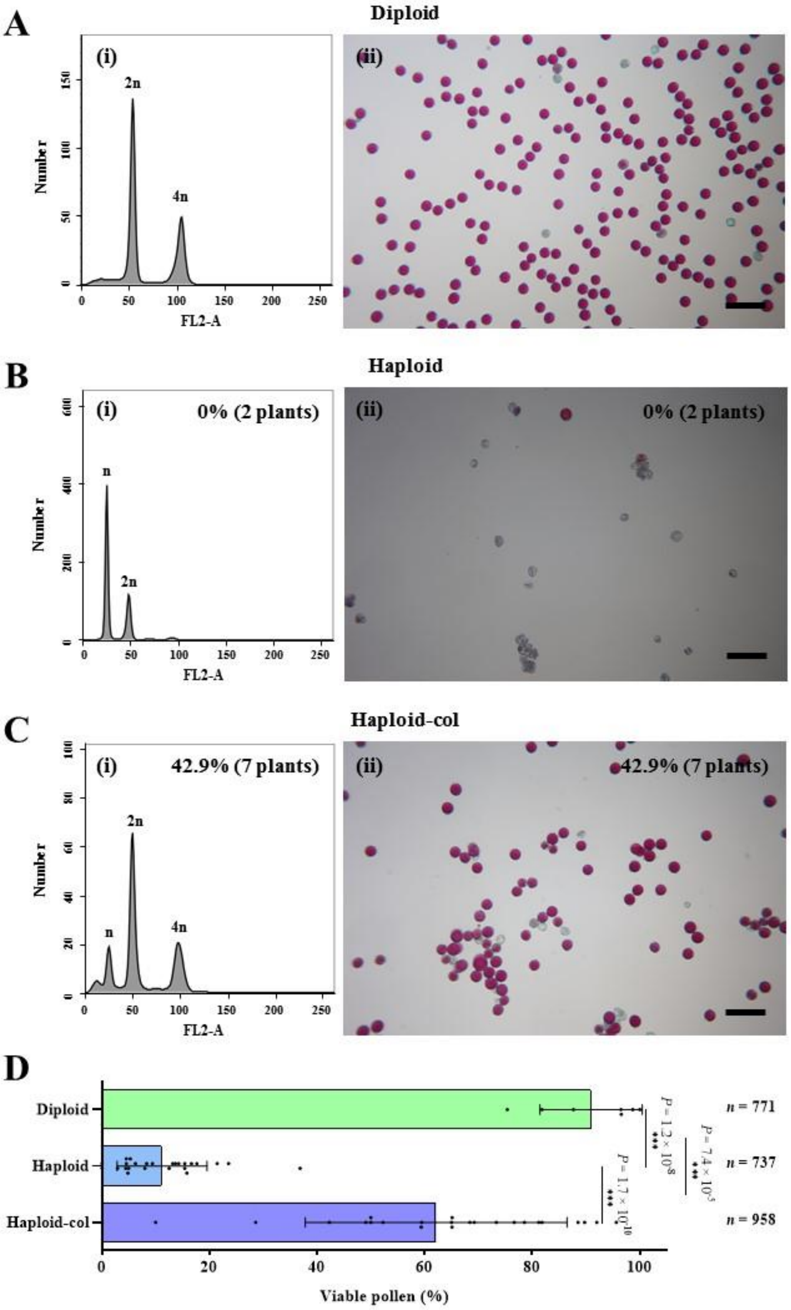


**Supplemental Figure 12. Colchicine induces somatic chromosome doubling in tomato haploids.** (**A** to **C**) Representative images of flow cytometry graphs (i) and pollen stained with Alexander's stain (ii) of diploid (A), untreated haploid (B) and haploid plant treated with colchicine (haploid-col) (C). Pollen that stained red were scored as viable. The percentages (shown in the upper right corner of each panels in (B) and (C)) are the ratio of plants showed the diploid cells (i) and high percentage of viable pollen (ii). Scale bar, 100 µm. (**D**) Quantification of pollen viability, shown as the percentage of viable pollen. Data represent the mean ± s.d.; ***$p < 0.001$ (two-tailed Student's $t$-test); $n$, the number of pollen grains.

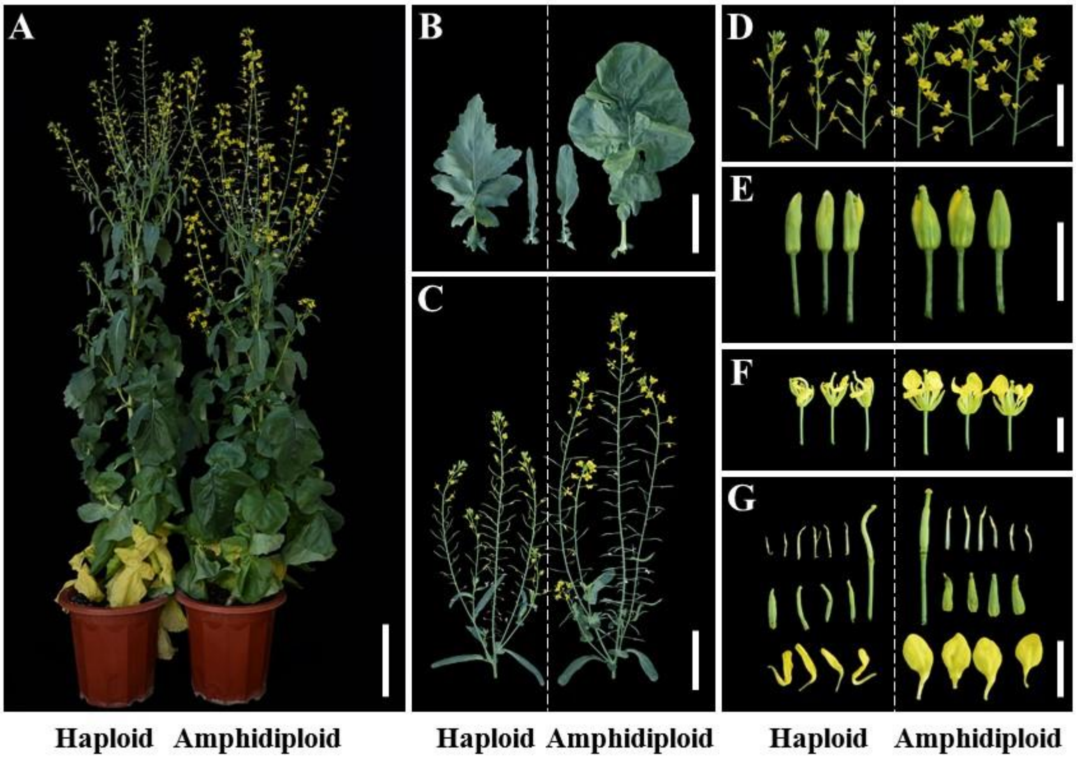


**Supplemental Figure 13. Phenotypes of rapeseed haploid plants.** Representative images of haploid and amphidiploid plants (**A**), leaves (**B**), inflorescences (**C** and **D**), flower buds (**E** and **F)** and dissected flower parts (**G**) of haploid and amphidiploid plants. Haploid plants have smaller vegetative and reproductive organs than amphidiploid plants. Scale bars: 20 cm (A), 10cm (B and C), 5 cm (D) and 1 cm (E to G).

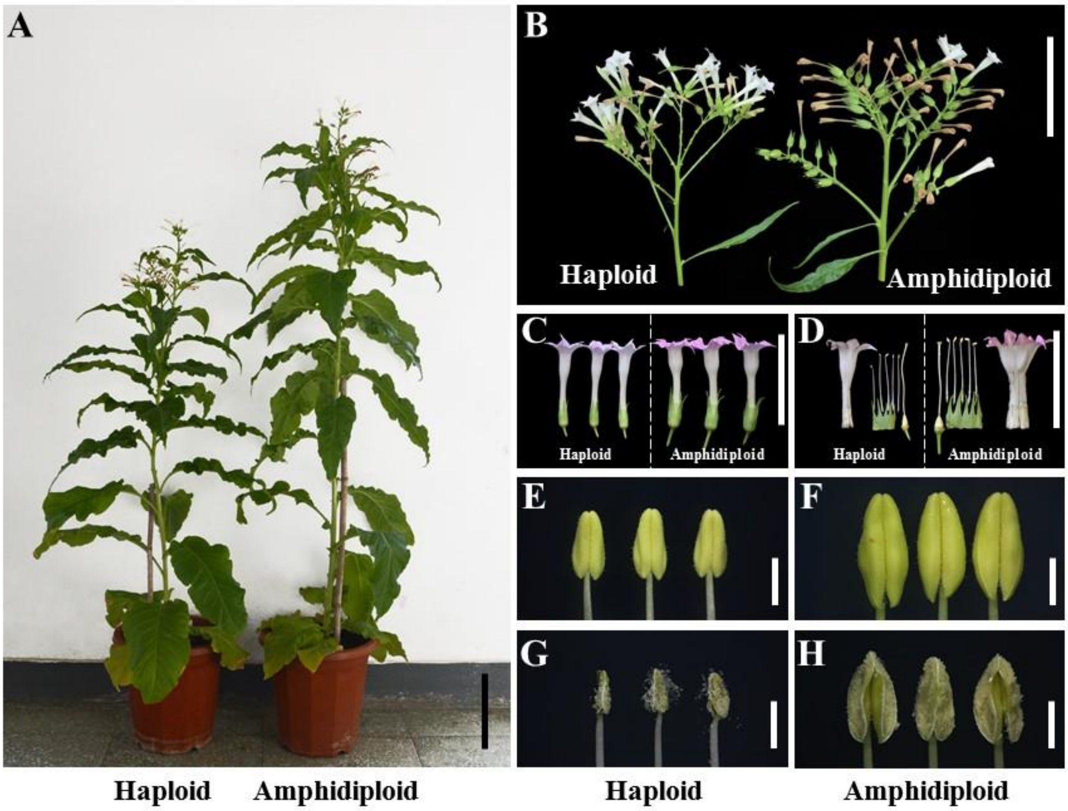


**Supplemental Figure 14. Phenotypes of tobacco haploid plants.** Representative images of haploid and amphidiploid plants (**A**), inflorescences (**B**), flower (**C)**, dissected flower parts (**D**) and mature anthers before (**E** and **F**) and after (**G** and **H**) dehiscence. Haploid plants have smaller vegetative and reproductive organs than tetraploid plant. Scale bars: 20 cm (A), 10cm (B), 5 cm (C) and 2 mm (E to H).

**Supplemental Table 1.** ***DMP*** **genes in selected dicot crops**

| Species | Family | Gene ID | Domain | Number of TM | Expression pattern |
|---|---|---|---|---|---|
| *Brassica napus* | *Brassicaceae* | *BnDMP1A* (*BnaA03g55920D*) | DUF679 | 4 | Anther, stamen |
| *Brassica napus* | *Brassicaceae* | *BnDMP1C* (*BnaC03g03890D*) | DUF679 | 4 | Anther, stamen |
| *Brassica napus* | *Brassicaceae* | *BnDMP2A* (*BnaA04g09480D*) | DUF679 | 4 | Anther, stamen |
| *Brassica napus* | *Brassicaceae* | *BnDMP2C* (*BnaC04g31700D*) | DUF679 | 2 | Anther, stamen |
| *Solanum lycopersicum* | *Solanaceae* | *SlDMP* (*Solyc05g007920*) | DUF679 | 4 | Pollen, flower bud |
| *Capsicum annuum* | *Solanaceae* | *CaDMP* (*T459_27262*) | DUF679 | 4 | NA |
| *Nicotiana tabacum* | *Solanaceae* | *NtDMP1* (LOC107762412) | DUF679 | 4 | NA |
| *Nicotiana tabacum* | *Solanaceae* | *NtDMP2* (LOC107783066) | DUF679 | 4 | NA |
| *Nicotiana tabacum* | *Solanaceae* | *NtDMP3* (LOC107807404) | DUF679 | 4 | NA |
| *Gossypium hirsutum* | *Malvaceae* | *GhDMP1* (LOC107911807) | DUF679 | 4 | Stamen |
| *Gossypium hirsutum* | *Malvaceae* | *GhDMP2* (LOC107924398) | DUF679 | 4 | Stamen |
| *Glycine max* | *Fabaceae* | *GmDMP1* (*GLYMA_18G097400*) | DUF679 | 3 | Flower bud |
| *Glycine max* | *Fabaceae* | *GmDMP2* (*GLYMA_18G098300*) | DUF679 | 3 | Flower bud |
| *Cucumis sativus* | *Cucurbitaceae* | *CsDMP* (*Csa_1G267250*) | DUF679 | 4 | Male flower bud |

DUF, domain of unknown function. TM, transmembrane. NA, not available

**Supplemental Table 5. Haploid induction rate using $T_1$ generation *sldmp* mutants.**

| Female parent | Male parent | Genotype | Seed setting rate (%) | Total plants | Haploids | HIR (%) |
|---|---|---|---|---|---|---|
| Ailsa Craig | Ailsa Craig | Wild type | 82.76 | 103 | 0 | 0.00 |
| AC34-T1-2 | AC34-T1-2 | *sldmp* | 22.49 | 55 | 1 | 1.82 |
| MT.AC | Ailsa Craig | Wild type | 81.97 | 869 | 0 | 0 |
| | AC12-T1-4 | *SlDMP/sldmp* | ND | 1,728 | 2 | 0.12 |
| | AC12-T1-19 | *sldmp* | ND | 507 | 1 | 0.20 |
| | AC34-T1-11 | *sldmp* | 26.41 | 160 | 1 | 0.63 |
| | MT9-T1-3 | *sldmp* | 21.15 | 143 | 1 | 0.70 |
| DF199 | Ailsa Craig | Wild type | 72.04 | 298 | 0 | 0 |
| | AC12-T1-19 | *sldmp* | ND | 38 | 1 | 2.63 |
| M.M82 | Ailsa Craig | Wild type | ND | 678 | 0 | 0.00 |
| | AC12-T1-4 | *SlDMP/sldmp* | ND | 530 | 2 | 0.38 |
| AF01 | AC12-T1-29 | *sldmp* | ND | 125 | 1 | 0.80 |
| | AC12-T1-39 | *sldmp* | ND | 591 | 2 | 0.34 |
| M.VF36 | AC12-T1-4 | *SlDMP/sldmp* | ND | 1,631 | 4 | 0.25 |
| Moneymaker | AC12-T1-4 | *SlDMP/sldmp* | ND | 421 | 1 | 0.24 |
| DF1 | MT9-T1-7 | *sldmp* | ND | 71 | 1 | 1.41 |
| Micro-Tom | MB-T1-Mix | *sldmp* | 5.51 | 262 | 2 | 0.76 |

The female parents are described in Supplemental Table 4. The *SlDMP* alleles of each male parent are listed in Supplemental Table 3. MB-T1-Mix, pollen was a mix of different T1 *dmp* mutant lines in the Moneyberg background. Haploids were first screened by molecular markers and then confirmed by flow cytometry. The haploid induction rate (HIR) was calculated with the formula: HIR (%) = (number of haploids) / (total plants) × 100%. ND, not determined.

**Supplemental Table 7. Accuracy of tomato haploid identification based on the FAST-Red marker.**

| Female parent | Male parent | Total seeds | White seeds | Weak RFP | Putative haploid | Haploids | Accuracy rate (%) | False negative rate (%) |
|---|---|---|---|---|---|---|---|---|
| DF199 | AC34-T2-Mix | 1,943 | 57 | 28 | 28 | 28 | 100 | 0 (1,131) |
| MA | AC34-T2-Mix | 8,513 | 151 | 117 | 117 | 117 | 100 | 0 (1,172) |
| DM | AC34-T2-Mix | 2,079 | 82 | 49 | 49 | 49 | 100 | NA |
| JZ801 | AC34-T2-Mix | 1,346 | 25 | 24 | 24 | 24 | 100 | NA |

The female parents are described in Supplemental Table 4. AC34-T2-Mix, haploids derived from different AC34-T2 lines. The number in the brackets indicates the number of red seeds screened. Accuracy rate (%) = (number of haploids) / (putative haploid number) × 100%. False negative rate (%) = (number of haploids from red seeds) / (number of red seeds) × 100%.

**Supplemental Table 8. Spontaneous chromosome doubling of tomato haploid plants.**

| Haploid plant ID | Pedigree | Pollen | Pollen germination | Fruit set | Seed set |
|---|---|---|---|---|---|
| AC34-T2-39 | AC34-T1-2 selfed | Yes | Yes | 4 | ND |
| Haploid 1 | MA × MT9-T1-3 | Yes | Yes | 3 | 4 |
| Haploid 2 | MA × AC12-T1-4 | Yes | Yes | 3 | ND |
| Haploid 3 | DF199 × AC12-T1-19 | Yes | Yes | 0 | ND |
| Haploid 4 | AF01 × AC12-T1-39 | Yes | Yes | 11 | 4 |
| Haploid 5 | AF01 × AC12-T1-29 | Yes | Yes | 6 | ND |
| Haploid 6 | DF1 × MT9-T1-7 | Yes | Yes | 8 | ND |
| Haploid 7 | AF01 × AC12-T1-39 | Yes | Yes | 0 | ND |
| Haploid 8 | MM × AC12-T1-4 | Yes | Yes | 8 | 19 |

ND, not determined.
